# *In vivo* and *In vitro* Characterization of the ClpC AAA+ ATPase of *Chlamydia trachomatis*

**DOI:** 10.1101/2022.08.22.504891

**Authors:** Stefan Pan, Aaron A. Jensen, Nicholas A. Wood, Beate Henrichfreise, Heike Brötz-Oesterhelt, Derek J. Fisher, Peter Sass, Scot P. Ouellette

**Affiliations:** Department of Microbial Bioactive Compounds, Interfaculty Institute of Microbiology and Infection Medicine, University of Tuebingen, Germany; Department of Pathology and Microbiology, College of Medicine, University of Nebraska Medical Center, Omaha, Nebraska, USA; Institute for Pharmaceutical Microbiology, University of Bonn, Meckenheimer Allee 168, 53115 Bonn, Germany; Cluster of Excellence-Controlling Microbes to Fight Infections, Tuebingen, Germany; School of Biological Sciences, Southern Illinois University Carbondale, Carbondale, Illinois, USA

**Keywords:** Chlamydia, ClpC, ClpP, AAA+ ATPase, Clp protease, differentiation, development

## Abstract

Bacterial AAA+ unfoldases are crucial for bacterial physiology by recognizing specific substrates and, typically, unfolding them for degradation by a proteolytic component. The <u>c</u>aseinolytic <u>p</u>rotease (Clp) system is one example where a hexameric unfoldase (e.g., ClpC) interacts with the tetradecameric proteolytic core ClpP. Unfoldases can have both ClpP-dependent and ClpP-independent roles in protein homeostasis, development, virulence, and cell differentiation. ClpC is an unfoldase predominantly found in Gram-positive bacteria and mycobacteria. Intriguingly, the obligate intracellular Gram-negative pathogen *Chlamydia*, an organism with a highly reduced genome, also encodes a ClpC ortholog, implying an important function for ClpC in chlamydial physiology. Here, we used a combination of *in vitro* and *in vivo* approaches to gain insight into the function of chlamydial ClpC. ClpC exhibits intrinsic ATPase and chaperone activities, with a primary role for the Walker B motif in the first nucleotide binding domain (NBD1). Furthermore, ClpC binds ClpP1P2 complexes via ClpP2 to form the functional protease ClpCP2P1 *in vitro*, which degraded arginine-phosphorylated β-casein. *In vivo* experiments confirmed that higher order complexes of ClpC are present in chlamydial cells. Importantly, the *in vivo* data further revealed severe negative effects of both overexpression and depletion of ClpC in *Chlamydia* as revealed by a significant reduction in chlamydial growth. Here again, NBD1 was critical for ClpC function. Hence, we provide the first mechanistic insight into the molecular and cellular function of chlamydial ClpC, which supports its essentiality in *Chlamydia*. ClpC is, therefore, a potential novel target for the development of anti-chlamydial agents.

**Significance:** *Chlamydia trachomatis* is an obligate intracellular pathogen and the world’s leading cause of preventable infectious blindness and bacterial sexually transmitted infections. Due to the high prevalence of chlamydial infections along with negative effects of current broad-spectrum treatment strategies, new anti-chlamydial agents with novel targets are desperately needed. In this context, bacterial Clp proteases have emerged as promising new antibiotic targets, since they often play central roles in bacterial physiology and, for some bacterial species, are even essential for survival. Here, we report on the chlamydial AAA+ unfoldase ClpC, its functional reconstitution and characterization, individually and as part of the ClpCP2P1 protease, and establish an essential role for ClpC in chlamydial growth and intracellular development, thereby identifying ClpC as a potential target for anti-chlamydial compounds.

## Introduction

Differentiation from one form to another in any organism is a complex process requiring the careful coordination of regulatory factors acting at each level of gene expression and protein homeostasis. In bacteria, the caseinolytic protease (Clp) system is known to contribute to regulated proteolysis and has been well studied in this regard [1-4]. The bacterial Clp protease is a highly conserved macromolecular protease comprised of two functionally distinct components: a tetradecameric proteolytic ClpP core and a corresponding hexameric type I AAA+ (<u>A</u>TPases <u>a</u>ssociated with diverse cellular <u>a</u>ctivities) unfoldase such as ClpX or ClpC [5]. Regulated protein hydrolysis requires the interaction of ATP-fueled AAA+ unfoldases with the ClpP core. The unfoldase recognizes, unfolds, and translocates protein substrates into the degradation chamber of the ClpP complex [5, 6]. Importantly, AAA+ unfoldases also function as protein chaperones independently of ClpP [7-10] and thus are crucial for bacterial physiology via multiple routes. These include both ClpP-dependent and ClpP-independent roles in protein homeostasis, development, virulence, cell differentiation, genetic competence, and, in some instances, viability [11-15]. Unsurprisingly, in recent years, ClpP and AAA+ unfoldases have emerged as promising target structures for drug development [16-23].

*Chlamydia trachomatis* (Ctr) is the world’s leading cause of preventable infectious blindness and bacterial sexually transmitted infections. In 2018, the CDC estimated that the prevalence of chlamydial infections was around 2.4 million within the United States, with associated healthcare costs imposing an economic burden of $691 million dollars [24]. However, the true infection rate and healthcare costs are likely underestimated, as approximately 70% of the cases are asymptomatic [25]. Current treatment modalities rely on broad-spectrum antibiotics that can have negative impacts on normal microflora within the genital tract [26]. Hence, there is an urgent need to develop, in the short-term, better anti-chlamydial treatment strategies by employing novel targets and mechanisms of antibiotic action and, in the long-term, vaccines capable of preventing infection. Regarding the former, we and others have demonstrated the utility of targeting chlamydial proteases to eradicate infection in a cell culture model [27-30]. Interestingly, in terms of new anti-chlamydial strategies, compounds that were derived from known ClpP-targeting dysregulators were highly effective at preventing chlamydial growth, although via a yet unknown, potentially ClpP-unrelated mechanism, while simultaneously having a limited impact on a variety of other bacterial species [31]. Taken together, it emerges that targeting the chlamydial Clp protease complex could be a novel strategy for anti-chlamydial treatment. A more comprehensive understanding of the individual Clp components in *Chlamydia* will facilitate this goal.

*Chlamydia* species are obligate intracellular Gram-negative bacteria that undergo a unique and complex biphasic developmental cycle to propagate. Infection of a susceptible host cell begins with the internalization of the small, electron dense, and non-dividing form of *Chlamydia* called the elementary body (EB) [32, 33]. Once internalized into a host-derived vacuole termed the inclusion, the EB immediately begins a primary differentiation event into the larger, less electron dense, non-infectious replicating form termed the reticulate body (RB). The first cell division is followed by rapid multiplication of RBs by an asymmetric polarized cell division mechanism [34, 35]. Secondary differentiation proceeds asynchronously as RBs condense into EBs [34]. These EBs will then exit the host cell through host cell lysis or extrusion of the inclusion itself. Given the distinct functional, morphologic, and proteomic differences between EBs and RBs, as well as the processes of differentiation that generate them, we have hypothesized that protein turnover is a critical and essential aspect of *Chlamydia* physiology and pathogenesis.

In evolving to obligate intracellular dependence, *Chlamydia* species have significantly reduced their genome size and content, and this is notable for *C. trachomatis*, which encodes roughly 900 open reading frames (ORFs) in ∼1Mbp. This suggests that the presence of a given gene in the genome indicates an important, if not essential, function. Of interest, *C. trachomatis* encodes two *clpP* genes (*clpP1* and *clpP2*). Of further interest is the presence of two AAA+ unfoldases encoded within the chlamydial genome: *clpX* and *clpC*. The *clpP2* and *clpX* genes are organized within an operon while *clpC* and *clpP1* are positioned separately on the chromosome [27, 28]. That *Chlamydia* should maintain two copies of a gene in its chromosome suggests an important function, especially since bacteria with two or more *clpP* genes are comparatively rare. We previously investigated the ClpX and ClpP protease components of *C. trachomatis in vitro* [28, 36], revealing the presence of an atypical, mixed ClpP core containing both ClpP1 and ClpP2. Combined with the AAA+ unfoldase ClpX, the chlamydial ClpXP complex displayed robust proteolytic activity for model substrates both *in vivo* [36] and *in vitro* [28, 36]. We have further demonstrated that overexpressing catalytic mutant, but not wild-type, isoforms of ClpP1, ClpP2, or ClpX *in vivo* in *C. trachomatis* has a negative impact on chlamydial growth and development [27]. These data support our hypothesis that the Clp system is critical for developmental cycle progression in *Chlamydia*.

While ClpX is well conserved across all types of eubacteria and even mitochondria, ClpC is typically found in Gram-positive bacteria and mycobacteria. ClpC has been shown to recognize substrates containing a phosphorylated arginine (via the activity of the McsB kinase) and is associated with the type III heat shock response in *B. subtilis* [37]. Given the presence of this second unfoldase, not typically found in Gram-negative bacteria, we hypothesized that ClpC may serve a specialized and essential function in *Chlamydia*. The current study was designed to characterize the *in vitro* enzymatic activity of this second AAA+ unfoldase, ClpC, and its putative interplay with ClpP in regulated proteolysis as well as the *in vivo* effects of dysregulating ClpC expression in *Chlamydia*. Multiple sequence alignments of chlamydial ClpC between heterologous species demonstrated that the chlamydial ortholog possesses the critical residues necessary for activity, including two nucleotide binding domains (NBD1/2). Principle findings from our study indicate that the chlamydial ClpC functions as an ATPase capable of delivering a substrate to the ClpP1P2 protease core for degradation. Further, we demonstrated the detrimental effects on chlamydial growth of dysregulating ClpC expression through exogenous overexpression or knockdown of *clpC* transcripts. Overall, our efforts have expanded our understanding of the function of the Clp protease system in *Chlamydia* by revealing that the activity of ClpC is essential for normal developmental cycle progression.

## Results

### Chlamydial ClpC retains the conserved motifs essential for ClpC activity

The conserved Walker-type nucleotide binding domains (NBDs) represent hallmark features of the Hsp100 protein family and are required for ATPase activity [38, 39]. Unlike ClpX, an Hsp100 chaperone with a single NBD (i.e., class II), ClpC contains two highly conserved NBDs (NBD1, NBD2; i.e., class I). To examine if *C. trachomatis* ClpC has the expected sequence elements to function as a class I AAA+ unfoldase, we took a bioinformatics approach by aligning its primary amino acid sequence to those of well-studied ClpC orthologs from model bacteria. Conserved regions and functional motifs are indicated in Figure 1A. The two canonical Walker A/B motifs of the NBDs characteristic of ClpC proteins are highly conserved between orthologous proteins, suggesting ATPase function is conserved in the chlamydial protein (red, purple). Importantly, the critical glutamate residues involved in ATP hydrolysis are present in the Walker B motif (E306/E644 in the chlamydial protein) [40]. The middle domain contact sites (yellow) necessary for oligomerization are present, albeit with an alteration in the chlamydial ortholog from phenylalanine 462 to tyrosine (F462Y). The IGF domain (green), important for interactions with ClpP, is also present. The N-terminal domain involved in the recognition of phosphorylated arginine (pArg) and other specific substrates and adaptors [41, 42] is also present in the chlamydial ClpC, but, perhaps not surprisingly given the likely substrate differences between species, shows more sequence variability beyond the conserved pArg recognition residues. Unique to the chlamydial ClpC is a ∼20 amino acid serine-rich linker connecting the N-terminal substrate recognition domain to the ATPase domain.

**Figure 1:**
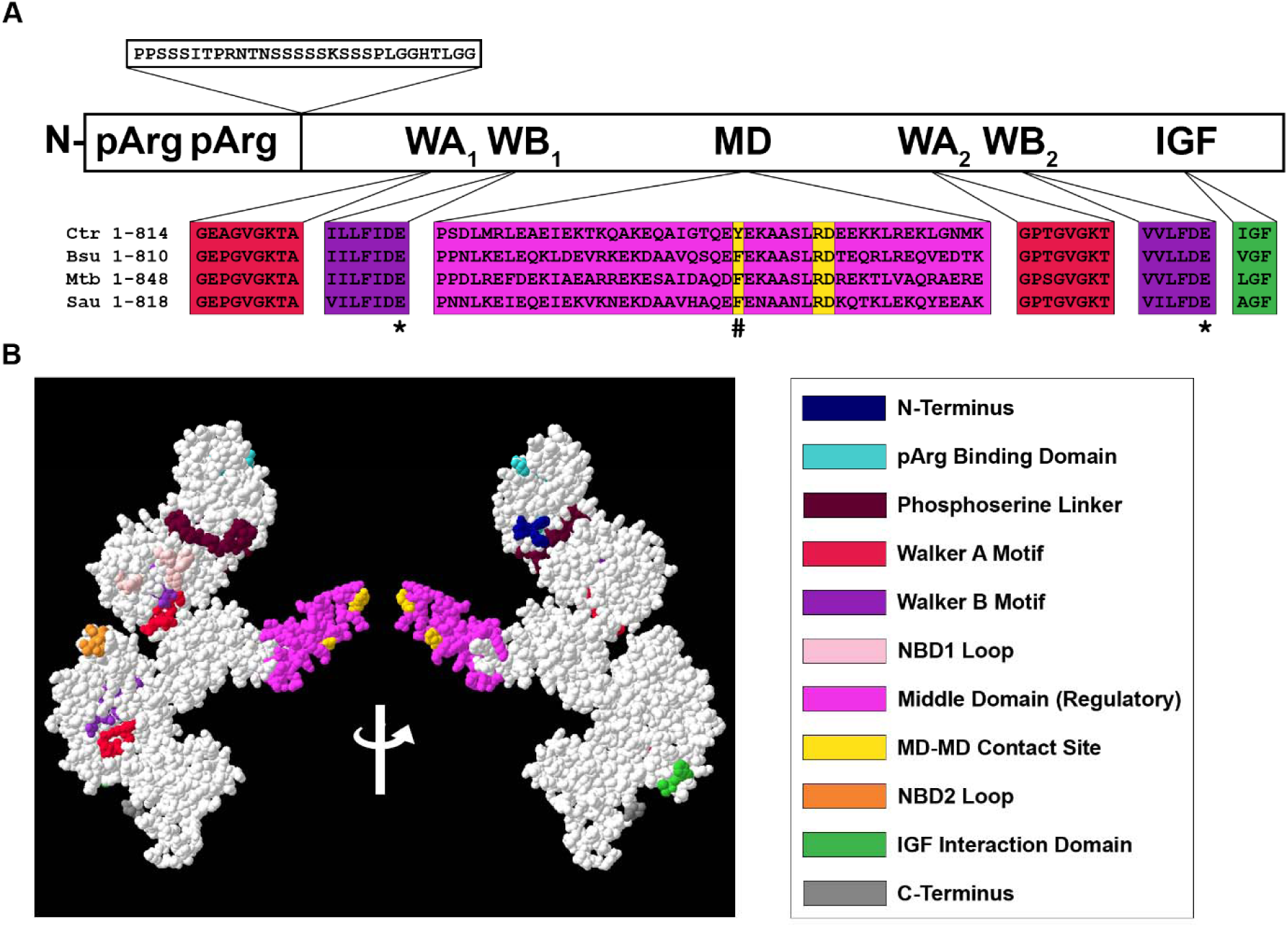
Chlamydial ClpC retains conserved functional domains as compared to ClpC orthologs. (A) Multiple sequence alignment of ClpC amino acids between Gram-negative *C. trachomatis* (Bu 434) and Gram-positive *B. subtilis* (strain 168), *M. tuberculosis* (H37Rv), and *S. aureus* (PS47). Color code is shown below (D1 or D2, domain 1 or 2, respectively; NBD, nucleotide binding domain; MD-MD, middle-domain middle-domain contact site for oligomerization; IGF Interaction Domain, region of interaction with ClpP binding cleft). The mutated glutamic acid sites within the Walker B motifs are denoted by *. The non-conserved tyrosine within the MD-MD contact site is denoted by #. Sequences were taken from Uniprot, aligned using Clustal Omega, and edited on Jalview v2.11.1.4. (B) 3-D color-coded modeling of ClpC monomer viewed from each face (navy, N-terminus; grey, C-terminus). Phyre2 was used to perform an intensive sequence analysis where the protein data bank file was visualized using Swiss-PdbViewer v4.1.0 for 3D modeling.

We subsequently modeled the chlamydial ClpC using a protein databank from Phyre2 and 3D-rendered using SWISS-MODEL software. As seen in Figure 1B, the 3D model of ClpC shows the Walker A/B motifs within close proximity to bind and hydrolyze ATP. The IGF domain is located at a surface exposed region consistent with where ClpP would bind. Overall, these data indicate a high likelihood that the chlamydial ClpC ortholog is a *bona fide* ClpC ATPase.

### Clustering analysis indicates chlamydial ClpC and ClpX are highly conserved across species

In homology relationship analyses, we previously showed that the ClpP isoforms of *C. trachomatis* are non-paralogous as both ClpP1 and ClpP2 were located at distinct and separate subgroups [28]. In the current study, we extended and included the amino acid sequences of the AAA+ unfoldases ClpX and ClpC into a homology relationship plot based on CLANS (CLuster ANalysis of Sequences) [28, 43], thereby obtaining a comprehensive picture of homology relationships of the entire chlamydial Clp system. In addition to 597 unique ClpP sequences from the UniProtKB/Swiss-Prot database, 493 unique ClpX sequences from the UniProtKB/Swiss-Prot database and 1671 unique ClpC sequences (34 from UniProtKB/Swiss-Prot and 1637 from UniProtKB/TrEMBL database) were used to generate a cluster map based on all-against-all pairwise sequence similarities (Fig. 2A). In agreement with our previous results, ClpP1 and ClpP2 are spatially separated from each other, despite their apparent interdependence on a functional level [27, 28]. For chlamydial ClpX and ClpC, respectively, nearly all sequences appeared to form one single homogenous cluster and, in contrast to ClpP, no distinct sub-cluster formation was computable. Also, the highly conserved (L/I/V)-G-(F/L) ClpP-binding motif, which exists in both chlamydial ClpC and ClpX, was found in 84% of all ClpC and 89% of all ClpX sequences despite high sequence variability in non-conserved regions (Fig. 2B), thus indicating a close paralogous relationship between ClpX/C orthologs and a highly conserved local protein domain architecture.

**Figure 2:**
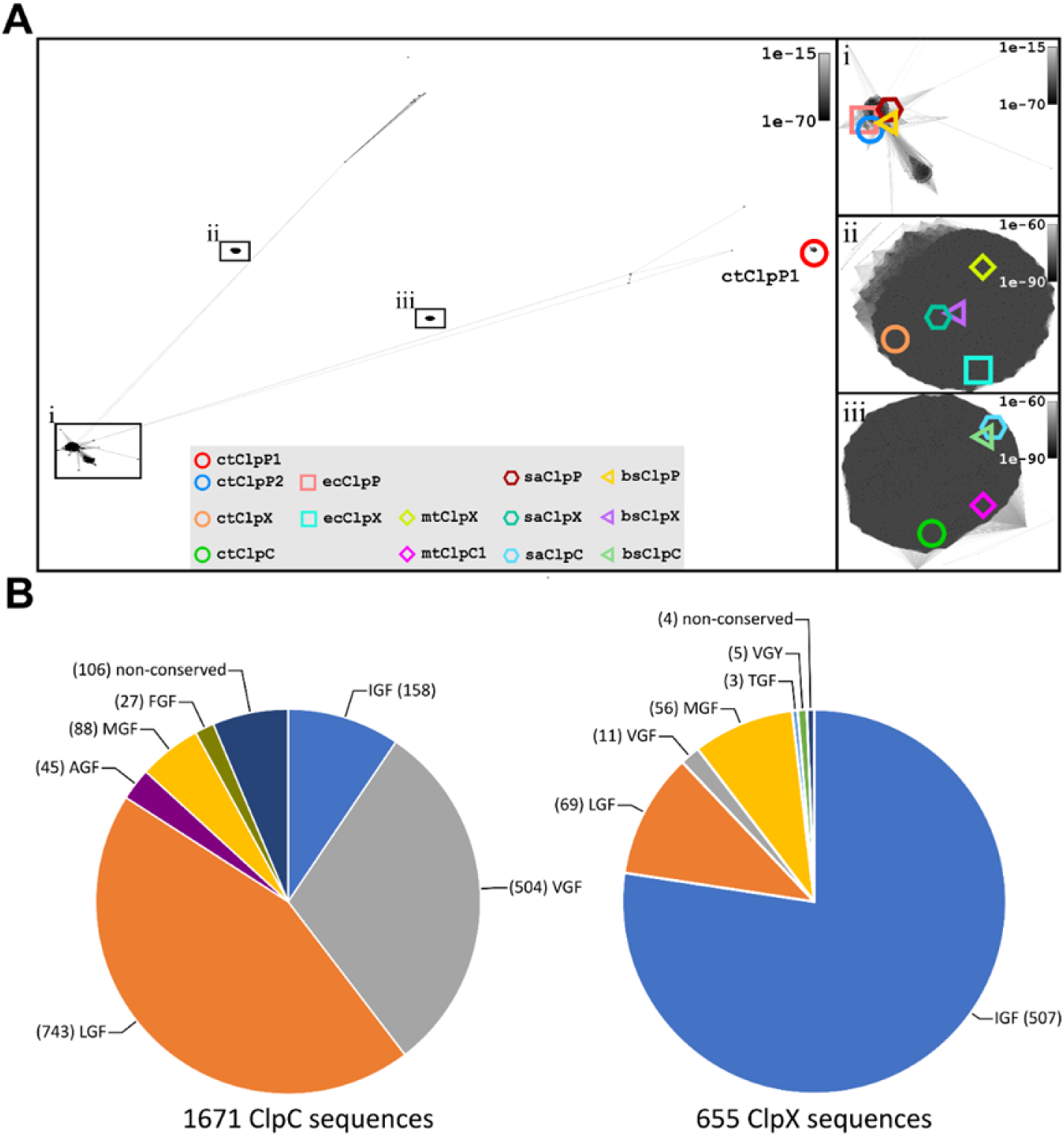
Cluster analyses of ClpP, ClpX and ClpC. (A) Distance plots based on CLANS (CLuster ANalysis of Sequences) [43]. P-value threshold was set to 1e-75. (i) Enlarged view of the major ClpP clusters. Most bacterial ClpP sequences (437/597) such as from *E. coli, S. aureus*, and *B. subtilis* form one major supercluster that also includes ctClpP2, although located in a distinct subgroup of the supercluster, but not ctClpP1. (ii) Enlarged view of the major ClpX cluster. All input ClpX sequences (493/493) densely cluster in a single group, no distinct subcluster formation was detected. (iii) Enlarged view of the major ClpC cluster. Almost all input ClpC sequences (1629/1637) densely cluster in a single group. For ClpC and ClpX, the range of extracted pairwise attraction values (based on corresponding HSP P-values) were too narrow to allow for subcluster visualization. (B) Hydrophobic pocket binding motif conservation in ClpC and ClpX. Typical variants of the ClpP-binding motif of (L/I/V)-G-(F/L) were found in 84% of all ClpC and 89% of all ClpX sequences. Partially conserved binding motifs such as (A/F/M/T/Y)-GF are found in roughly 10% of each Clp-ATPases. Remaining sequences (6% for ClpC, 0.6% for ClpX) showed no apparent conservation.

### The Walker B motif in the first nucleotide binding domain of ClpC disproportionally contributes to ATPase and chaperone activities

To begin characterizing chlamydial ClpC, we investigated its ATPase activity *in vitro* using recombinant protein preparations. In a quantitative ADP to ATP conversion assay coupled to luciferase activity readout, wild-type ClpC exhibited ATPase activity in a dose-dependent manner (Fig. 3A). Furthermore, inactivation of the Walker B motif in NBD1, conferred by introducing an E306A mutation, disproportionally inhibited ATP hydrolysis as compared to the inactivation of the Walker B motif in NBD2 (E644A). While inactivation of NBD2 reduced ATPase activity by approximately 30%, mutation of NBD1 reduced overall ATPase activity by over 70%, indicating a dominant role of the NBD1 Walker B motif for ATPase activity of ClpC.

**Figure 3:**
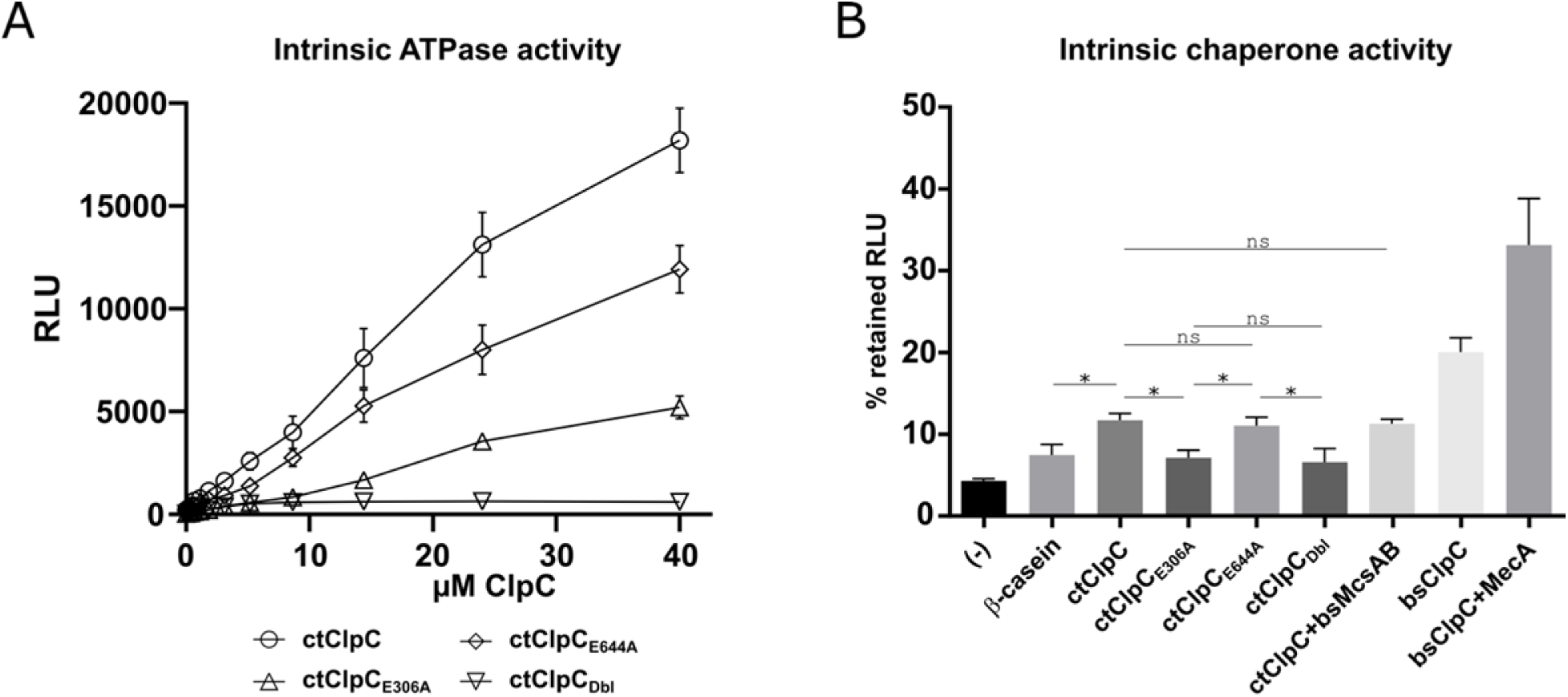
ATPase and chaperone activities of chlamydial wild-type and mutant ClpC. (A) Intrinsic ATPase activity of wild-type and mutant ClpC. When using a quantitative ADP to ATP system followed by luciferase activity readout, wild-type ClpC displayed ATPase activity in a dose-dependent manner. An E306A mutation in the Walker B motif of NBD1 strongly impaired overall ATPase activity. In contrast, ATPase activity was less affected by mutation of the Walker B motif in NBD2 (E644A). ATPase activity was entirely lost in the Walker B double mutant (ctClpC_Dbl_). (B) Chaperone activity of wild-type and mutant ClpC. In a protein aggregation prevention assay, the residual activity of heat-treated luciferase was determined in the absence or presence of either β-casein (non-chaperone control assay) or ClpC from *C. trachomatis* (ct) or *B. subtilis* (bs) as indicated. In the absence of β-casein and ClpC (-), heat-treated luciferase showed a 96% reduction of its initial activity and thus served as the minimal luminescence baseline. β-casein was used as reference protein to assess non-specific protein-stabilizing effects. Wild-type ctClpC demonstrated low but significant chaperone activity, indicated by an increase in the retained luciferase activity compared to the β-casein control. In contrast, the ctClpC_E306A_ and ctClpC_Dbl_ mutants were devoid of chaperone activity, since luciferase activity was similar to the β-casein control, whereas the ctClpC_E644A_ mutant showed wildtype-like chaperone activity, thus corroborating the importance of the Walker B motif in NBD1 for ClpC activity. The statistical significance analysis was performed via two-way ANOVA determination (*p<0.05; ns: not significant). As a positive control, chaperone activity was also determined for bsClpC, which was substantial and could be further elevated by the adaptor protein MecA.

Next, we analyzed the chaperone activity of Ctr ClpC in a protein aggregation prevention assay using heat-induced denaturation of luciferase. Compared to the control assay using β- casein, the addition of wild-type ClpC from either *C. trachomatis* or *B. subtilis* (Bs) increased luciferase activity and thus prevented to some degree the heat-induced denaturation of the luciferase protein (Fig. 3B). Corroborating our results for ClpC ATPase activity, the Walker B motifs in NBD1 and NBD2 displayed an unequal enzymatic contribution to the chaperone activity. While the ClpC_E644A_ mutant (inactivated Walker B motif in NBD2) showed unaltered chaperone activity compared to wild-type ClpC, the ClpC_E306A_ mutant (inactivated Walker B motif in NBD1) and the double mutant ClpC_Dbl_ (E306A, E644A) did not prevent luciferase aggregation above the β-casein control, thereby highlighting the importance of the Walker B motif in NBD1 for ClpC activity. Addition of β-casein did lead to a low level of stabilization of the luciferase as indicated by an increase of luciferase activity above background where neither β-casein nor wild-type ClpC was added. This suggests some nonspecific stabilizing effects by proteins with non-chaperone functions, which is supported by the E306A and Dbl mutant ClpC. Of note, the addition of *B. subtilis* McsAB kinase (bsMcsAB), which confers arginine phosphorylation of substrates (Fig. S1), did not enhance Ctr ClpC chaperone activity, suggesting arginine phosphorylation may not be a prerequisite for protein disaggregation via ClpC under the conditions tested.

### ClpC binds to the heteromeric ClpP1P2 complex via ClpP2 to form the functional protease ClpCP2P1

Access to the catalytic sites of ClpP is non-permissible for polypeptides and proteins, since these are commonly too large to freely diffuse through the axial pores into the inner lumen of the ClpP core, thereby restricting substrate hydrolysis by the ClpP core to small peptides up to 30 residues [5, 44-47]. To assess whether ClpC could activate ClpP protease activity *in vitro*, we first prepared arginine-phosphorylated β-casein (casein-pArg) as a protein substrate for ClpCP [41, 48]. Successful substrate phosphorylation was confirmed by pArg-specific antibodies (Fig. S1). In subsequent protease activity assays, we then analyzed the proteolytic capacity of different combinations of ClpC, ClpP1, and ClpP2 to digest casein-pArg (Fig. 4). Here, ClpP1P2 did not notably digest casein-pArg in the absence of ClpC. Similarly, ClpC alone did not affect the abundance of casein-pArg. Also, incubation of either ClpP1 or ClpP2 in the presence of ClpC was not sufficient for proteolytic activity. However, when ClpP1, ClpP2, and ClpC were combined, casein-pArg was clearly degraded, thereby showing that ClpC-mediated proteolytic activity of the chlamydial Clp protease depends on the presence of a heteromeric complex consisting of both ClpP1 and ClpP2. These *in vitro* protease assays indicate that, like ClpX [28], ClpC also forms a functional protease with ClpP1P2.

**Figure 4:**
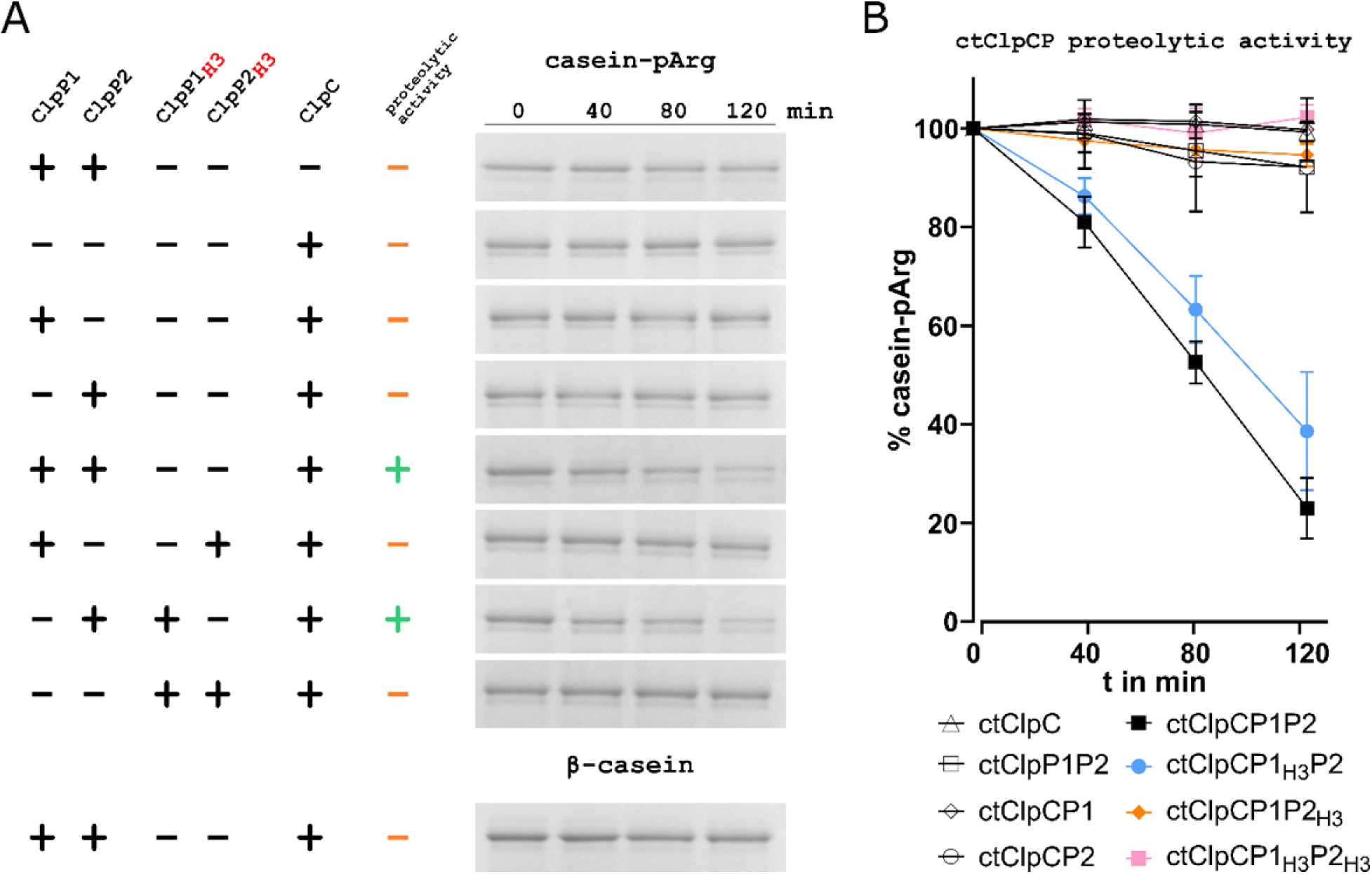
Proteolytic activity of the ClpCP complex. (A) Protease activity as determined by degradation of arginine-phosphorylated β-casein (casein-pArg) as a substrate and assessed using SDS-PAGE. Individually, neither ClpP1 nor ClpP2 were sufficient to degrade casein-pArg regardless of the presence of ClpC. Casein-pArg degradation was only observed when ClpP1, ClpP2, and ClpC were present, supporting that ClpC interacts with a heteromeric ClpP1P2 core. To determine the binding partner of ClpC, ClpP1_H3_ and ClpP2_H3_ mutant proteins were constructed that carry triple amino acid exchanges in the Clp-ATPase-binding regions. While the degradation of casein-pArg was unaltered in the presence of ClpP1_H3_P2 and ClpC, proteolytic activity was clearly inhibited when ClpP1P2_H3_ or ClpP1_H3_P2_H3_ were used together with ClpC, indicating an interaction of ClpC with the ClpP1P2 core via ClpP2. Of note, as a control, unphosphorylated β-casein was not degraded in the presence of ClpCP2P1, indicating the requirement of arginine-phosphorylation for ClpC-mediated degradation. (B) Densitometry of bands shown in Figure 4A. Experiments were performed in triplicate, and mean values are presented.

We next sought to identify the binding partner of ClpC within the heteromeric ClpP core. Unlike *clpX*, which is encoded within an operon with *clpP2*, the *clpC* gene is not located closely to either *clpP1* or *clpP2* [27, 28], and thus interaction of ClpC with either ClpP homolog was not predictable based on the genomic context. To characterize ClpC binding to ClpP, we constructed mutants of ClpP1 and ClpP2 with amino acid changes in their respective Clp ATPase binding regions based on known homologous ClpP sequences (i.e., ClpP1_H3_ (V57A, V77A, L186T) and ClpP2_H3_ (F63A, F83A, I190T)). When we substituted wild-type ClpP1 and/or ClpP2 for the respective mutant proteins in casein-pArg degradation assays, ClpCP2_H3_P1 as well as ClpCP2_H3_P1_H3_ were devoid of proteolytic activity, while casein-pArg degradation was almost fully retained in the ClpCP2P1_H3_ mutant (Fig. 4). Of note, unphosphorylated β-casein was not degraded by ClpCP2P1, indicating the requirement of arginine-phosphorylation for ClpC-mediated degradation in these assays. Our results thus support a model where ClpC binds to the heteromeric ClpP1P2 complex via ClpP2.

### Both wild-type and Walker B mutant isoforms of ClpC oligomerize *in vivo*

A key characteristic of ClpC orthologs necessary for their activity is their ability to form a homohexamer through a trimer of dimers [49]. Although we detected *in vitro* enzymatic activity of the recombinant ClpC protein, we wanted to verify that higher order complexes were present *in vivo*. Therefore, to determine whether ClpC oligomerizes, we performed a series of *in vivo* experiments using a bacterial two-hybrid system (BACTH) and crosslinking followed by western blotting during infection with wild-type *C. trachomatis* serovar L2. We also tested oligomerization of isoforms with mutations in the Walker B motifs (i.e., E306A, E644A, and both mutations (Dbl)) to assess and confirm that these mutations did not impair the ability of the isoform to form oligomers in other analyses.

The BACTH system operates by the reconstitution of adenylate cyclase activity when the two catalytic fragments, T25 and T18 of the *Bordetella pertussis* Cya, are expressed as fusion proteins with proteins of interest and are brought into close proximity by the interacting proteins [50]. When expressed separately or as fusions with non-interacting proteins, no cAMP is produced. Reconstituted cAMP production activates β-galactosidase expression, which can be qualitatively assessed on 5-bromo-4-chloro-3-indolyl-β-D-galactopyranoside (X-Gal) plates and quantitatively measured by enzyme assays. For BACTH assays, we cloned the wild-type or mutant *clpC* genes into the pKT25 or pUT18C vectors, which were then co-transformed into a Δ*cya Escherichia coli* strain (DHT1). As expected, we observed both positive qualitative and quantitative results when assessing homotypic interactions (i.e., wild-type with wild-type) (Fig. 5A&B). Heterotypic interactions between wild-type and mutants were also analyzed, and these resulted in positive interactions as well (Fig. 5A&B). Importantly, we did not observe interactions between ClpC and ClpX, which served as a negative control for these studies. These results demonstrate the ability of ClpC isoforms to oligomerize both homotypically and heterotypically. Therefore, we expect that ectopically expressed ClpC isoforms can potentially form mixed complexes with endogenous ClpC in *Chlamydia*. This is consistent with previous data for the other chlamydial Clp components and is important for subsequent data interpretation [36].

**Figure 5:**
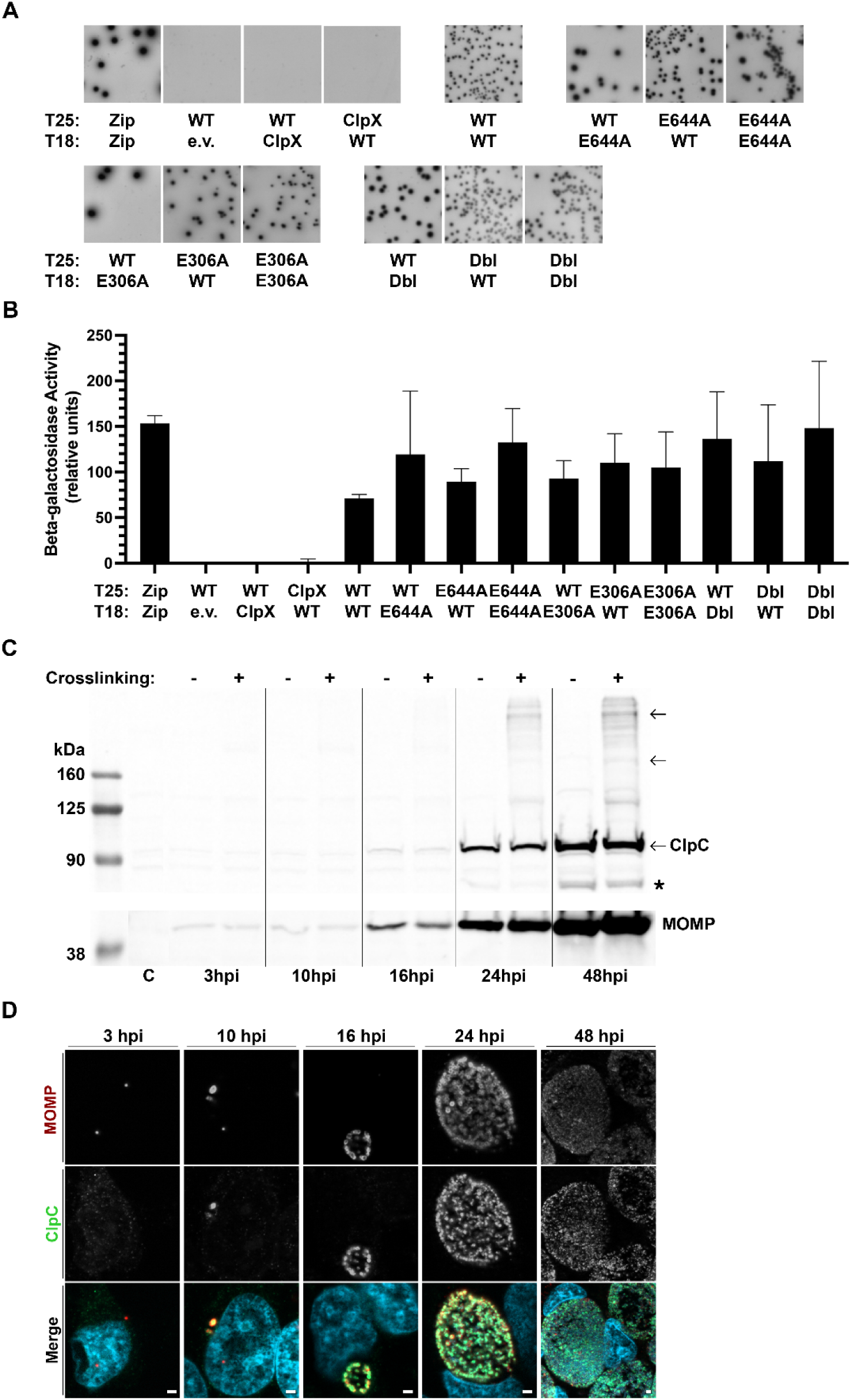
Wild-type and mutant ClpC isoforms interact both homotypically and heterotypically as assessed by *in vivo* bacterial two-hybrid assay and crosslinking of endogenous ClpC in cell culture. (A) Bacterial adenylate cyclase two-hybrid (BACTH) assays showing homotypic interactions between wild-type (WT) or mutant (E306A, E644A, or Dbl) ClpC isoforms, and heterotypic interactions between wild-type and mutant ClpC isoforms. Positive (Zip:Zip) or negative (T25-WT_ClpC versus T18-empty vector, and WT_ClpC versus Ctr ClpX unfoldase in either orientation) controls are also shown. Photos have been set to grayscale. (B) Quantification of β- galactosidase activity of the BACTH interactions shown in panel (A) in arbitrary units. Positive interactions are considered as having at least five times the activity of the negative controls. (C) *In vivo* assessment of endogenous ClpC oligomerization. Wild-type Ctr L2 infected HeLa cells were crosslinked or not at the indicated times post-infection with the primary amine crosslinker disuccinimidyl suberate (DSS). DMSO was used as vehicle control in non-crosslinked samples with protease inhibitors added with DSS before incubation. 25 micrograms of protein from cell lysates were added per lane for SDS-PAGE prior to transfer for western blotting. Arrows indicate expected ClpC monomer, dimer, and hexamer sizes. A degradation product of ClpC within protein samples was consistently observed and is denoted by *. Uninfected HeLa cell lysate was included as a negative control and denoted as C. (D) Detection of endogenous ClpC by immunofluorescence. Antibody staining of wild-type Ctr L2 of samples collected in parallel with (C), fixed with MeOH at the indicated times post-infection, and stained for ClpC (green), major outer membrane protein (MOMP, bacteria; red), and DNA (blue). Merged images of all color channels are shown. Representative images were taken in triplicate on a Zeiss Apotome at ×100 magnification with ×5.5 digital zoom for 3-24 hpi and ×2.75 digital zoom for 48 hpi. Scale bar = 2 µm.

To confirm oligomerization during infection, we infected HeLa cells with wild-type *C. trachomatis* L2 and collected protein lysates at varying times post-infection from samples that had been crosslinked or not using disuccinimidyl suberate (DSS). 25 µg of whole cell lysates were separated by SDS-PAGE, transferred to PVDF membranes, and probed with an antibody for endogenous ClpC (Fig. S2). Western blot data indicated the presence of a ClpC monomer of the expected size (95 kDa) in all conditions starting at 16 hours post-infection (hpi) (Fig. 5C), consistent with our prior study using an alternatively-derived antibody [27]. Interestingly, in samples treated with DSS, higher order banding patterns were observed that were consistent with both dimers (190 kDa) and hexamers (570 kDa). Additional bands were present in the crosslinked samples, which may reflect potential substrates or other interaction partners. This is under investigation. Immunofluorescence analysis (IFA) of control samples revealed ClpC as early as 10 hpi, a time when transcripts for *clpC* are increasing (Fig. 5D) [27]. Overall, these data confirm that chlamydial ClpC can oligomerize *in vivo* to form the expected hexamer necessary for activity.

### Overexpression of wild-type ClpC is detrimental to *Chlamydia*

We have previously assessed the effects of overexpression of wild-type and mutant isoforms of ClpP1, ClpP2, and ClpX on chlamydial growth and viability [27, 36]. In general, overexpression of mutant but not wild-type Clp proteins was detrimental to *Chlamydia*. To determine the impact of overexpression of the ClpC isoforms, we created plasmids encoding anhydrotetracycline (aTc)-inducible, 6xHis-tagged constructs and transformed them into *C. trachomatis* L2 lacking its endogenous plasmid (-pL2). As a control, we also examined the effect of overexpressing mCherry from the same plasmid backbone. HeLa cells were infected with the different strains, ClpC or mCherry expression was induced at 4 hpi (Fig. 6A) or 10 hpi (Fig. S3), and cells were subsequently fixed for IFA to determine the effects on bacterial and inclusion morphology or harvested for inclusion forming unit (IFU) assays to quantify growth effects (Fig. 6B). As with endogenous ClpC, we confirmed that the ectopically expressed ClpC isoforms also formed higher order structures *in vivo* (Fig. S4).

**Figure 6:**
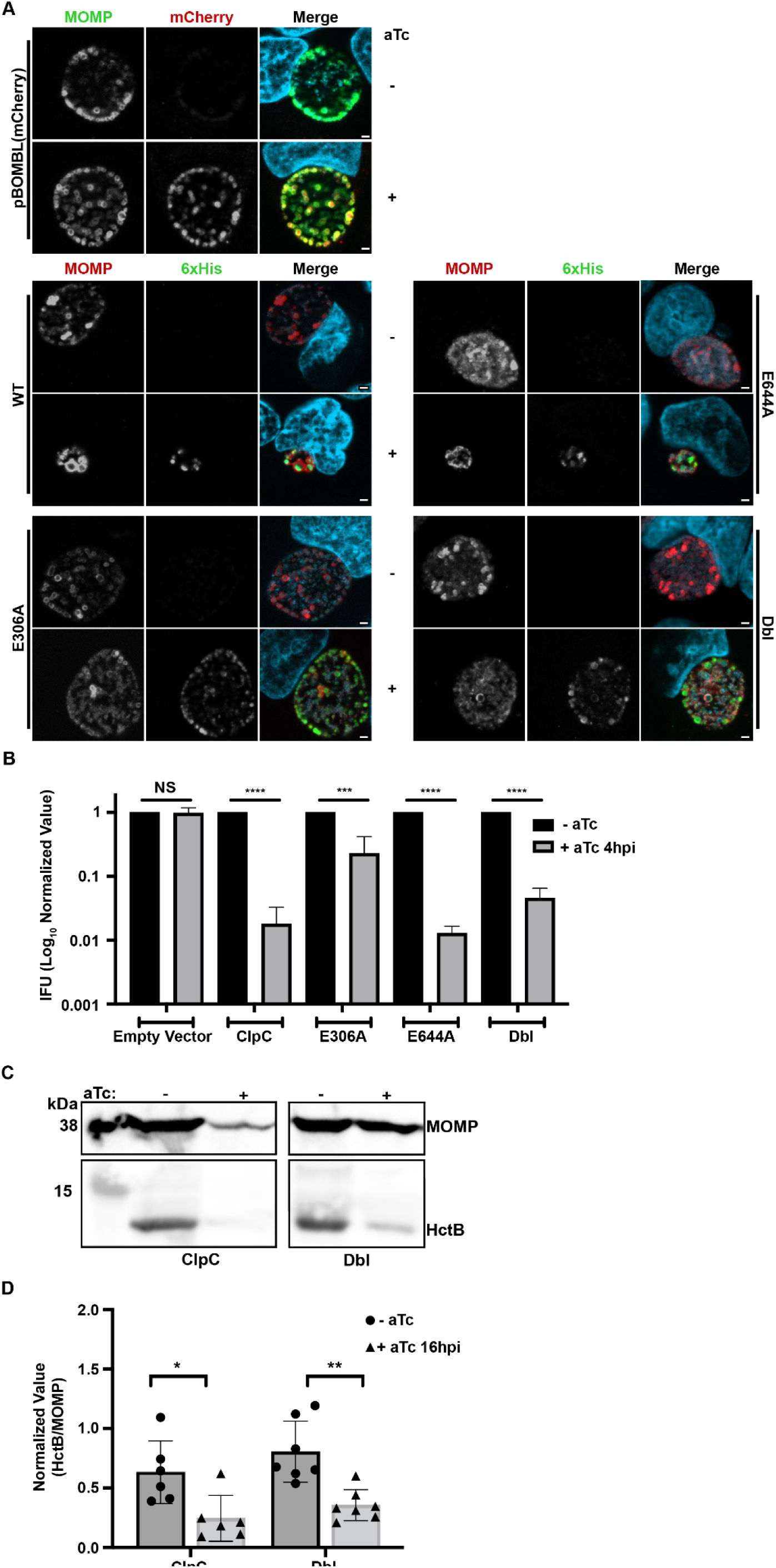
Overexpression of ClpC isoforms shows NBD1 is critical for function and detrimental to chlamydial growth. (A) Effect of overexpression of ClpC isoforms or mCherry on chlamydial inclusion size. Immunofluorescence analysis of the control plasmid expressing mCherry or exogenously overexpressed ClpC_6xHis isoforms as indicated (WT, E306A, E644A, Dbl (E306A/E644A)). Cells were infected with the transformants, and expression of the constructs was induced or not at 4 hpi with 20 nM aTc. At 24 hpi the control plasmid was fixed with 3.25% formaldehyde and 0.125% glutaraldehyde followed by MeOH permeabilization to retain mCherry fluorescence, while the ClpC isoforms were fixed with MeOH. Samples were stained for MOMP (green for mCherry strain/red for ClpC strains), or 6xHis (green) tagged protein from plasmid induction, and DNA (light blue). Representative images were taken in triplicate on a Zeiss Apotome at ×100 magnification with ×5.5 digital zoom. Scale bar = 2 µm. Note the small size of the inclusions in the ClpC WT and E644A +aTc panels. (B) Quantification of chlamydial growth after expressing or not the ClpC isoforms or mCherry. Recovered IFUs of cells infected with each transformant from (A). Samples were collected in triplicate and combined at 24 hpi, frozen at −80°C, and titrated onto a fresh monolayer of cells to quantify inclusions in the secondary infection. Inclusions were identified by counting the GFP-positive inclusions. Data are the average of at least three independent experiments. Note the large decrease in growth when expressing the WT, E644A, and Dbl mutant isoforms as compared to mCherry or E306A. (C & D) Representative images (C) of 25 µg whole protein lysate ran on western blot and stained for the EB specific protein HctB with MOMP functioning as the loading control. Whole cell lysates were collected at 48 hpi from HeLa cells infected with Ctr exogenously expressing ClpC or Dbl induced at 16 hpi with 20 nM aTc. Note the relative differences in HctB staining to MOMP after inducing the ClpC isoforms. MOMP = 42 kDa, HctB = 13.9 kDa. (D) Densitometric analysis of (C), where HctB staining was normalized with MOMP. Data are the average of at least three independent experiments. The statistical significance analysis was performed via a parametric, unpaired Student’s T-test (*p<0.05; **p<0.01; ***p<0.001; ****p<0.0001).

For IFA, fixed cells were labelled with an anti-6xHis antibody to examine induced ClpC expression or directly assessed for mCherry fluorescence (control plasmid), an anti-major outer membrane protein (MOMP) antibody as a marker for individual bacteria, and DAPI to stain DNA. There was no observable impact of overexpressing mCherry on inclusion size or bacterial numbers (Fig. 6A). In contrast to what we observed with the other Clp proteins [27, 36], overexpression of wild-type ClpC resulted in a stark difference between uninduced and induced samples. Whereas typical inclusion sizes were observed in the uninduced conditions, overexpression of wild-type ClpC resulted in noticeably smaller inclusions with fewer organisms. A similar phenotype was observed when overexpressing the E644A mutant (inactivated Walker B motif in NBD2). However, corroborating our *in vitro* results, the inclusion morphology was not obviously altered for the E306A mutant (inactivated Walker B motif in NBD1) or for the Dbl mutant. Results from the IFU assay (equivalent to a CFU assay for most bacteria) were generally consistent with the IFA data (Fig. 6B). Overexpression of wild-type ClpC and the E644A mutant resulted in a significant, approximately 100-fold decrease in IFU production. Interestingly, the Dbl mutant displayed a significant reduction (∼30-fold) in IFUs even though the IFA appeared to be unaffected compared to the uninduced samples whereas overexpression of the E306A mutant resulted in a statistically significant decrease in growth (∼5-fold) that was much less than the other isoforms. These results indicate (i) that *Chlamydia* is highly sensitive to increased wild-type or mutant ClpC levels and (ii) that, consistent with our *in vitro* activity assays, the NBD1 is more important for ClpC activity *in vivo*.

To further explore why the wild-type and Dbl mutant overexpression strains displayed similar IFU recoveries but disparate morphological effects by IFA, we examined whether secondary differentiation, as assessed by the levels of the EB-associated protein HctB, showed differences in these strains. To this end, we infected cells and induced overexpression at 16 hpi and subsequently collected protein lysates at 48 hpi from both strains and their uninduced controls. We induced at 16 hpi to ensure adequate chlamydial protein in the wild-type strain given the dramatic inhibition in inclusion growth and development when overexpressing wild-type ClpC. Given the effects of wild-type ClpC overexpression on bacterial and inclusion growth, the levels of HctB normalized to MOMP during overexpression were significantly reduced compared to the uninduced control, as expected (Fig. 6C&D). Similarly, and consistent with the IFU data if not the IFA data, the normalized levels of HctB were also reduced in the Dbl mutant when overexpressed (Fig. 6C&D). These data indicate that overexpression of the wild-type ClpC isoform blocks developmental cycle progression (as reflected by low MOMP and low HctB) whereas overexpression of the Dbl mutant ClpC isoform prevents secondary differentiation while having a limited impact on overall inclusion growth and bacterial replication (i.e., high MOMP, low HctB).

Since both ClpC and ClpX interact *in vitro* with the same ClpP ortholog in *Chlamydia*, we wanted to ensure that the effects of ClpC overexpression *in vivo* were not a result of titrating ClpP away from ClpX. To test this, we co-expressed both wild-type ClpC with a 6xHis tag and wild-type ClpX with a FLAG tag as a transcriptional fusion. We have previously shown that overexpression of ClpX, like ClpP2, has limited effects on bacterial growth or inclusion morphology [27, 36]. Thus, any negative effect on inclusion morphology when co-expressing both ClpC and ClpX could be attributed to an effect of ClpC, assuming an equal affinity for the ClpP components. Cells were infected, and expression of the proteins was induced at 4 or 10 hpi before fixation for IFA or collection of samples for IFU assays at 24 hpi. Data from both IFA and IFU assays indicated that there was still a dramatic effect on inclusion morphology and a significant reduction in chlamydial growth (Fig. S5). These results indicate that the negative impact of ClpC overexpression is not likely due to ClpC’s titrating ClpP away from ClpX.

### Inhibition of *clpC* expression by CRISPR interference is detrimental to chlamydial growth

As *Chlamydia* was sensitive to increased levels of ClpC, we next wanted to examine the effects of reduced ClpC levels on chlamydial growth using an inducible dCas12 CRISPR interference (CRISPRi) knockdown system developed in our lab [51]. As *clpC* is transcribed during RB growth and division, and transcript levels peak at 14-16 hpi [27], we first validated that knockdown was occurring after dCas12 expression by collecting nucleic acid samples at 14 hpi, after inducing samples at 10 hpi (Fig. 7A). RT-qPCR analysis revealed an approximate 75% reduction in *clpC* transcripts, confirming knockdown in the *clpC*-targeting strain, but no significant effects on other control transcripts (i.e., *euo, incA,* and *clpP1*). There was no effect of dCas12 expression alone on any of the transcripts we assessed (Fig. 7A). Interestingly, we did not see any changes to inclusion or bacterial morphology by IFA at 24 hpi (after inducing dCas12 expression at 4 hpi) as compared to an empty vector control expressing dCas12 alone (Fig. 7B). We next examined the effects of *clpC* knockdown by IFU assay at 24 hpi by counting GFP-positive inclusions since GFP is constitutively expressed from the CRISPRi plasmid (Fig. 7C). Under these conditions, we measured a significant decrease in IFUs in samples where *clpC* transcripts had been knocked down. In addition, there was a significant reduction in overall plasmid retention after *clpC* knockdown (Fig. 7D), indicating that *Chlamydia* is highly sensitive to reductions in ClpC levels and loses the CRISPRi plasmid to become susceptible to penicillin, which is non-bacteriocidal in *Chlamydia* [52, 53]. Importantly, overexpression of dCas12 alone had only a limited effect on chlamydial growth and plasmid retention (Fig. 7C&D), as previously observed [51]. Overall, these data, taken together with the overexpression data, indicate alterations in ClpC levels and/or activity severely disrupt chlamydial growth and development.

**Figure 7:**
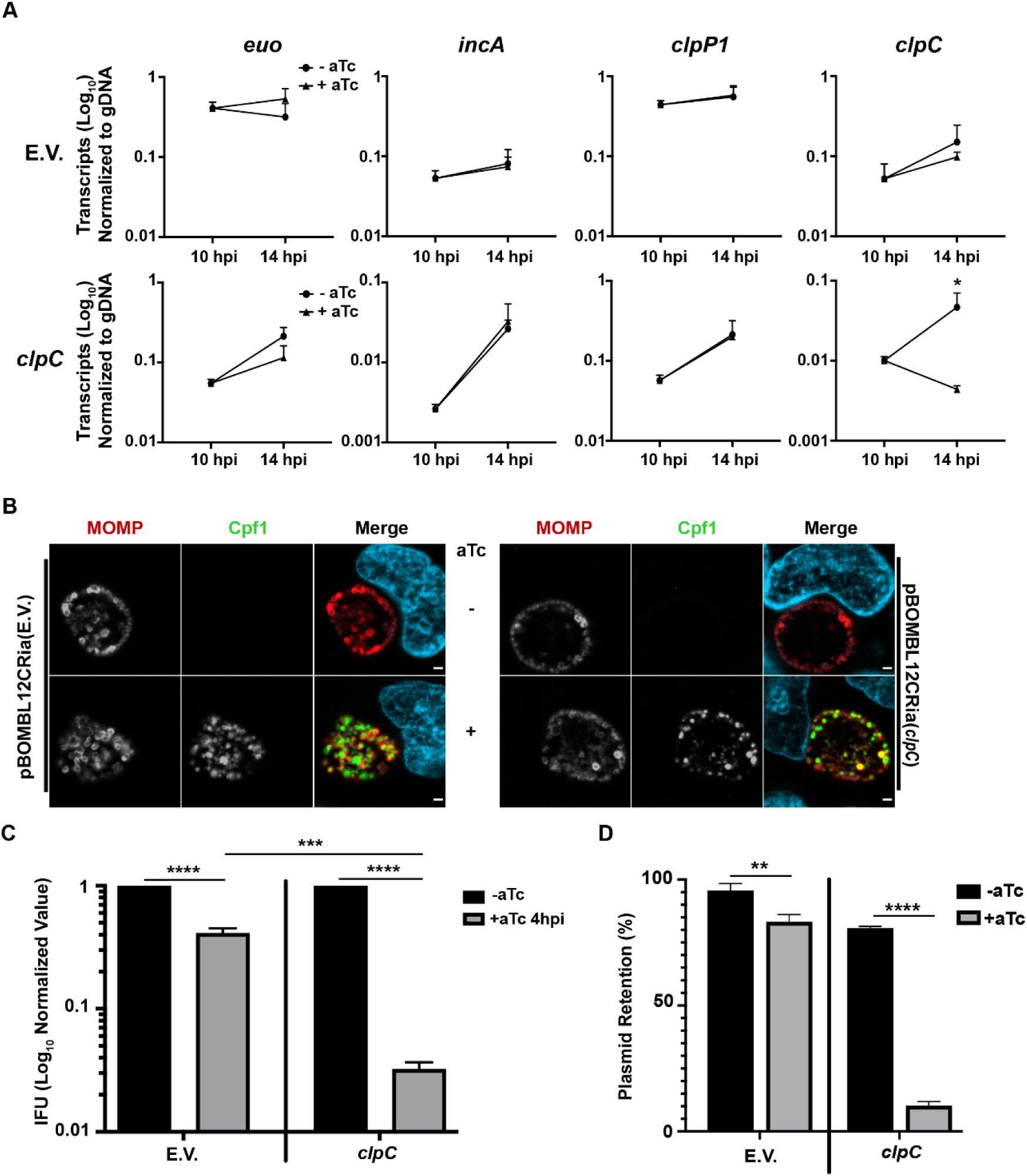
Knockdown of ClpC expression using CRISPR interference is detrimental to *C. trachomatis*. (A) Measure of *clpC* and control transcripts in strains expressing a *clpC*-targeting CRISPRi or dCas12 alone (i.e., empty vector). RT-qPCR of the indicated transcripts between uninduced and induced samples normalized to genomic DNA (gDNA). (B) Immunofluorescence analysis of the effects of knocking down, or not, *clpC* expression on bacterial morphology and inclusion size. IFA of the empty vector (e.v.) and *clpC* knockdown strain where samples were induced at 4 hpi with 2 nM aTc and fixed at 24 hpi with MeOH. Fixed cells were stained for MOMP (red), Cpf1 (for dCas12; green), and DNA (blue), then mounted on coverslips for imaging. Representative images were taken in triplicate on a Zeiss Apotome at ×100 magnification with ×5.5 digital zoom. Note the apparent absence of effect on inclusion sizes between the different strains and conditions. (C) Quantitative assessment of the effect of *clpC* knockdown on chlamydial EB production. IFU samples were collected in tandem with samples from (B) and quantified on a fresh cell layer as described in Materials and Methods. Note the large drop in IFUs when *clpC* transcripts are knocked down in spite of the lack of effect on inclusion size. (D) Measure of plasmid retention during knockdown of *clpC* and control conditions. Plasmid retention was examined as the ratio of GFP-positive (marker for the transformation plasmid) to MOMP-positive (total bacteria) inclusions in the IFU assay between uninduced and induced samples. The statistical significance analysis was performed via a parametric, unpaired Students T-test (*p<0.05; ***p<0.001; ****p<0.0001).

## Discussion

Prokaryotes express a variety of ATP-dependent protease complexes, such as the Clp and Lon proteases, that can perform regulated protein degradation [54-56]. For the Clp system, AAA+ unfoldases are critical for recognition and unfolding of substrates into the proteolytic chamber comprised of the barrel-shaped ClpP tetradecamers [5]. One such unfoldase is ClpC, a class I unfoldase with two nucleotide binding domains (NBD). ClpC is commonly found in Gram-positive bacteria. In organisms like *Bacillus subtilis*, ClpC is encoded within an operon containing accessory factors McsA/B that support ClpC substrate targeting by phosphorylating arginine residues on target proteins [55, 57]. ClpC recognizes and binds these phosphoarginines to deliver the substrate to ClpP. In many Gram-positive bacteria, ClpC has been shown to regulate vital cellular processes including stress responses, genetic competence, cell physiology, differentiation, and even virulence [58]. That the obligate intracellular Gram-negative pathogen *Chlamydia*, an organism with a highly reduced genome, would maintain an ortholog to ClpC is intriguing. Based on the characterized functions of ClpC in other bacteria, we hypothesized that ClpC is an important factor for developmental cycle progression and differentiation in *Chlamydia*.

We initiated our studies by performing *in silico* analyses through a bioinformatics approach. Based on sequence alignments, all the key residues and domains associated with substrate recognition and ATPase activity are present in chlamydial ClpC and share high homology with other ClpC orthologs (Figures 1&2). Of note, the chlamydial ClpC contains a novel serine-rich linker region between the N-terminal substrate binding domain and the first ATPase domain. Whether this linker can be phosphorylated as a regulatory mechanism or simply provides more flexibility for substrate recognition requires further study. Nonetheless, the observed stringent conservation of domain organization, critical motifs for ATPase activity, and ClpP-binding motif suggests a conserved mechanism of action of both ClpC and ClpX in most bacteria, including *C. trachomatis*.

Using recombinant proteins of different isoforms of chlamydial ClpC, we confirmed that wild-type ClpC has an *in vitro* dose-dependent intrinsic ATPase activity that was abolished by mutating both Walker B NBDs. However, NBD1 contributed to the majority of the intrinsic ATPase activity: the single E306A mutant reduced the activity by ∼70% while mutation of the NBD2 (E644A) decreased ATPase activity by only 30%. Given that NBD1 domains are suggested to have a major role in substrate unfolding, this result is perhaps not surprising, but indeed, it contrasts with previous findings on *B. subtilis* ClpC where Walker B NBD2 contributed to the majority (85%) of the intrinsic ATPase activity [4, 59]. Chlamydial ClpC also possessed some chaperone-like activity as it reduced aggregation of heat-treated luciferase. Again, the NBD1 was more important for this activity. Interestingly, in our hands, the activity of chlamydial ClpC was roughly half that of the *B. subtilis* ortholog, and, unlike *B. subtilis* ClpC, its chaperone activity could not be enhanced by the addition of bsMcsAB, which was used as a surrogate for the chlamydial McsAB due to ongoing difficulties purifying the proteins. Therefore, our data suggest differences for the individual activity and function of ClpC in *Chlamydia* and *B. subtilis*, which may relate to the different roles of Walker B NBD1 and NBD2 in both enzymes and the unique biology of *Chlamydia*.

Similar to our previous observation for chlamydial ClpX [28], ClpC also formed an active protease complex with a mixed ClpP1P2 tetradecamer *in vitro*. No functional proteolytic activity was observed when ClpC was present with either ClpP1 or ClpP2, individually. Our data revealed that interaction of ClpC with the ClpP1P2 core is mediated via the ClpP2 protein, thereby identifying ClpP2 as the central docking partner for both AAA+ ATPases, ClpC and ClpX, in *Chlamydia*. In this asymmetric complex, ClpP1 contributes to enzymatic activity together with ClpP2 [28] and may further serve as a crucial structural element for the assembly of the proteolytically competent ClpP core, similar to mycobacterial Clp [60-63], which may provide an explanation for the apparent functional interdependence of chlamydial ClpP1 and ClpP2. This is further supported by cluster analyses which locate ClpP1 far outside the ClpP “super cluster” (Fig. 2A), indicating pronounced differences of ClpP1 to ClpP2 and other ClpP orthologs that may explain the lack of binding to ClpC and ClpX.

Interestingly, our *in vitro* protease assays also revealed that the standard β-casein substrate typically used in such assays was not degraded. Rather, an arginine-phosphorylated form of β-casein was necessary for degradation to be detected. As noted above, ClpC is typically encoded with McsAB in an operon [58, 64]. Intriguingly, *Chlamydia* also possesses McsAB orthologs that are encoded as an operon but separated from *clpC* in the chromosome. The *mcsAB* operon is flanked by tRNAs, suggesting that it may have been horizontally acquired. Further work is necessary to determine the function of these proteins and whether arginine phosphorylation is occurring during the chlamydial developmental cycle. One intriguing possibility under investigation in our lab is that phosphorylation of arginine residues on critical substrates leads to their degradation to trigger differentiation. Such a model is consistent with the described activity of the ClpC/McsAB system in other bacteria [57, 64]. However, to evaluate this in *Chlamydia*, we need to identify potential substrates recognized by the ClpC unfoldase.

We next assessed the *in vivo* function of ClpC during chlamydial growth and development. We first examined the oligomerization state of endogenous ClpC during the developmental cycle using crosslinkers to preserve higher order complexes since ClpC functions as a hexamer in other bacteria. With this approach, we observed that both dimers and larger complexes, consistent with a hexamer, were detected starting at 16 hpi, the earliest time at which we detected endogenous ClpC. Notably, the bulk of protein was in a monomeric form, suggesting that ClpC may require activation to form a complex *in vivo*. This would be consistent with other systems [7, 57]. We also detected additional bands in the crosslinked samples that may represent substrates bound to ClpC, but this requires further investigation and is a major goal moving forward.

To evaluate effects of altering ClpC levels in *Chlamydia*, we created transformants carrying plasmids with inducible constructs encoding wild-type or NBD1/2 mutant (separately and together) isoforms. Based on our prior studies of chlamydial ClpX and ClpP, we had anticipated that overexpressing the wild-type isoform of ClpC would have a negligible impact on chlamydial growth whereas the mutant isoforms would impair developmental cycle progression [27, 36]. Surprisingly, we observed the opposite: a drastic reduction in chlamydial growth when overexpressing the wild-type ClpC but less of an effect when overexpressing the double NBD1/2 Walker B (E306A/E644A) mutant. Interestingly, overexpression of the Dbl ClpC mutant appeared to have normal morphology but had reduced IFU yields, suggesting that production of viable EBs may be impaired as we noted for the ClpX Walker B mutant [36]. Further, our *in vivo* data confirmed the importance of the NBD1 as overexpressing the E306A mutant had only a slight negative impact on chlamydial growth whereas overexpressing the E644A mutant resulted in a similar phenotype as the wild-type ClpC overexpression. Given that both ClpX and ClpC interact with ClpP2 *in vitro* to form an active protease, one interpretation of these data is that higher levels of ClpC may outcompete ClpX for ClpP2 and that this is detrimental because ClpX might no longer accomplish its function. Nonetheless, co-expressing both unfoldases resulted in the same phenotype as expressing ClpC alone, suggesting that competition between unfoldases for ClpP binding does not explain the ClpC overexpression phenotype. Rather, ClpC binding of an essential substrate(s) may lead to its degradation or sequestration. However, we cannot exclude that the unfoldases may have differential binding affinities for the ClpP complex, which requires further investigation.

As a complementary approach to our overexpression studies, we also implemented a CRISPR interference strategy developed in our lab to inducibly repress *clpC* transcription [51]. We confirmed successful knockdown of *clpC* transcript levels at the peak phase of its expression during the developmental cycle. Our results indicate that *Chlamydia* is highly sensitive to reduced ClpC levels as we observed a significant loss of the plasmid encoding the CRISPRi system, which resulted in the susceptibility of the organism to the selection agent (penicillin). Not surprisingly given this observation, we determined that reduced ClpC levels resulted in decreased chlamydial growth. These data were similar to the effects of overexpression of the Dbl mutant ClpC isoform, which is also expected to disrupt ClpC-mediated degradation of its substrates. Collectively, these data suggest that the inability of ClpC (by its absence or impairment) to target specific substrate(s) for degradation by ClpP or for possible chaperone functions impairs growth and/or differentiation in *Chlamydia*. Further, this indicates that ClpC is likely essential to *Chlamydia* and is consistent with failure to obtain a *clpC*::intron mutant (D. Fisher, unpublished observation) and loss of a *clpC*::Tn mutant from a transposon mutant pool during serial passaging [65]. Overall, both the overexpression and knockdown data suggest that the activity of ClpC must be tightly regulated to ensure proper developmental cycle progression.

In sum, the current work has helped advance our understanding of the Clp system in an important obligate intracellular pathogen. This study has demonstrated (i) that chlamydial ClpC has the function and activity of a canonical ClpC with the ability to recognize a phosphoarginine-tagged substrate for subsequent degradation by ClpCP2P1, and (ii) that alterations to ClpC levels have dramatic negative impacts on chlamydial growth and viability. Taken together, this indicates ClpC is likely essential to *Chlamydia* and represents a potential target for development of antimicrobial compounds.

## Materials and Methods

### Strains and cell culture

Recombinant Clp proteins from *C. trachomatis* were expressed using *E. coli* BL21 (DE3) *dAPX-1* as previously described to prevent undesired co-purification of *E. coli* Clp proteins [28]. Recombinant McsA and McsB of *B. subtilis* (bsMcsAB) and McsB of *Geobacillus stearothermophilus* (gsMcsB) were expressed using *E. coli* strain *C43*. Unless stated otherwise, cells were cultivated in lysogeny broth (LB) supplemented with ampicillin (100 µg/ml) at 37 °C under vigorous agitation. The human epithelial HeLa cells were used for all studies involving cell culture (overexpression, gDNA, protein extractions, plaque purification, transformation). HeLa cells were routinely passaged in Dulbecco’s modified Eagle’s medium (DMEM; Gibco/Thermo Fisher) with 10% fetal bovine serum (FBS; Sigma, St. Louis, MO) and were verified to be mycoplasma free by a LookOut Mycoplasma PCR Detection kit (Sigma). *C. trachomatis* serovar L2 EBs (25667R) (kind gift of Dr. Ian Clarke, University of Southampton) that lacked endogenous pL2 plasmid were prepared and used for transformation [66].

### Bioinformatic analysis

Gene sequences of *Chlamydia trachomatis* (Bu 434) were obtained from STDGEN database (http://stdgen.northwestern.edu) or KEGG Genome Browser [67-69]. The RefSeq protein sequences for *Staphylococcus aureus* (PS47), *Mycobacterium tuberculosis* (H37Rv), and *B. subtilis* (strain 168) were acquired from the NCBI protein database (http://www.ncbi.nlm.nih.gov/proteins/). Pairwise protein alignments for ClpC to find sequence identity were performed using the NCBI Protein BLAST function (https://blast.ncbi.nlm.nih.gov/Blast.cgi) [70]. Multiple sequence alignments were performed using Clustal Omega [71] with default settings, and formatted for presentation using Jalview Version 2 [72]. Homology relationship plots based on CLANS (CLuster ANalysis of Sequences [43]) were computed using ClpX, ClpC, and ClpP amino acid sequences and the BLOSUM62 scoring matrix. Input included 597 unique ClpP and 493 unique ClpX sequences extracted from the UniProtKB/Swiss-Prot database (reviewed sequences). Input ClpC sequences consist of 34 unique sequences from the UniProtKB/Swiss-Prot database and 1637 unique sequences (800 to 950 aa in length) from the UniProtKB/TrEMBL (unreviewed) database. BLAST HSP E-values up to 1e-10 were extracted for calculating pairwise attraction values. For iterative cluster formation, the P-value threshold was set to 1e-75. To examine predicted three-dimensional (3D) structure, PBD files were acquired from the Phyre2 website (http://www.sbg.bio.ic.ac.uk/phyre2/html/page.cgi?id=index) [73]. The acquired PBD file was then 3D-modeled using SWISS-MODEL, available on the ExPASy server [74-76].

### Plasmid construction

A full list of primers used in this study is provided in Table S1 within the supplemental material. Cloning of the ClpC expression plasmid was achieved by ligating PCR-amplified *C. trachomatis clpC* (CT_286, *C. trachomatis* D/UW-3/CX, NC_000117.1) into the pET-11a vector, which allows for the expression of fusion proteins with a C-terminal Strep-tag, using the NEBuilder system (New England Biolabs; NEB). Suitable primers for PCR were calculated by the NEBuilder Assembly Tool 2.0 and are listed in Table S1. For the construction of expression plasmids encoding bsMcsA (BSU_00840) and bsMcsB (BSU_00850) of *B. subtilis 168* (NC_000964.3), PCR-amplified inserts were obtained by using the primers listed in Table S1. The *mcsB* gene of *G. stearothermophilus* (*gsmcsB*) was amplified from pET-21a-gsmcsB_his [77] for subcloning using the primers listed in Table S1. All resulting PCR fragments were cloned into a modified pET-11a vector featuring two *Bsa*I non-identical restriction sites as a universal, non-palindromic polylinker followed by a C-terminal Strep(II) tag sequence (pET-MP3) [28].

Site-directed mutagenesis was achieved by using the “QuickChange II site-directed mutagenesis” kit (Agilent) according to the manufacturers protocol. Previously constructed plasmids encoding the hydrophobic pocket mutant proteins ClpP1_L186T_ and ClpP2_I190T_ [28] were used as templates to introduce additional mutations V57A and V77A for ClpP1_L186T_ and F63A and F83A for ClpP2_I190T_. See Table S1 for primers used for site-directed mutagenesis. Resulting ClpP1 and ClpP2 hydrophobic pocket triple mutants were referred to as ClpP1_H3_ and ClpP2_H3_. In the same manner, construction of ClpC Walker B motif mutants (E306A, E644A) was performed using the primers listed in Table S1. The resulting ClpC Walker B motif single mutants were referred to as ClpC_E306A_ and ClpC_E644A_, whereas the ClpC Walker B motif double mutant was referred to as ClpC_Dbl_. Sequence identity of all resulting constructs was verified by Sanger sequencing (LGC genomics).

BACTH constructs made for homo- and heterotypic interaction analysis were created using the HiFi Cloning (NEB) protocol. NEBuilder assembly tool was used to generate primers (http://nebuilder.neb.com). One set of primers each were designed for PCR amplification to insert products into either pKT25 or pUT18C [50]. *C. trachomatis* L2 (Bu 434) genomic DNA was used as template for wild-type ClpC, and the pST25 Q5 mutagenesis plasmid (shown below) as template for ClpC mutant PCR amplification with primers. PCR amplified products were confirmed for length by agarose gel electrophoresis. BACTH plasmids were generated by taking PCR amplified product and inserting them into the backbone of pKT25 or pUT18C cut with BamHI and KpnI. These HiFi reactions were transformed into DH5αI^q^ *E. coli* (NEB), and isolated plasmid was verified by restriction enzyme digest and sequencing by Eurofins Genomics. The sequence verified plasmids were then used for interaction studies in BACTH (see below).

Constructs made for chlamydial transformation were created using the HiFi Cloning protocol. NEBuilder assembly tool was used to generate primers. Primers were designed to add a poly-histidine (6xHis) or FLAG tag to the gene of interest being inserted on the 3’ end of the overlap into the shuttle vector. *C. trachomatis* L2 (Bu 434) genomic DNA was used as template for PCR amplification with primers, and products were confirmed for length by agarose gel electrophoresis. Overexpression plasmids were generated by first taking PCR amplified products and inserting them into the backbone pBOMBL::L2 [51] cut with EagI and KpnI to remove the mCherry gene. pBOMBL::L2 is a derivative of pBOMB4-Tet::L2 (kind gift of T. Hackstadt, NIH [50]) with a modified weakened ribosomal binding site to reduce leaky expression and off target effects when not induced. These HiFi reactions were transformed into DH10β *E. coli*, and isolated plasmid was verified by restriction enzyme digest and sequencing by Eurofins Genomics. The sequence verified plasmids were transformed into *C. trachomatis* (see below).

To generate mutations of the ClpC Walker B motifs, Q5 mutagenesis (New England BioLabs) was used. Primers were designed encoding either the E306A or E644A mutation of the Walker B motifs for PCR linearization of the plasmids. The ClpC BACTH construct pST25-ClpC was used as the template for both the PCR amplifications, which were recircularized by a kinase-ligase-DpnI (KLD) reaction. These reactions were then transformed into DH5α *E. coli* for plasmid generation. The plasmids were isolated, and mutations and sequence were confirmed by Sanger sequencing (Eurofins Genomics) prior to use in the BACTH system. These plasmids served as the templates for PCR products to move mutant *clpC* alleles into pBOMBL::L2, pKT25, or pUT18C.

### Protein expression

Expression and purification of recombinant chlamydial ClpP1 and ClpP2 was carried out as previously described [28]. *E. coli* cultures containing the expression plasmids for either ctClpC, bsMcsA, bsMcsB or gsMcsB were grown until mid-log exponential phase (OD_600_ of 0.6 to 0.7). Upon induction of expression by adding 1 mM IPTG, *E. coli* cultures were further shaken at 18 °C overnight. The following purification steps were executed at 4 °C. After centrifugation, cell pellet was resuspended in prechilled buffer A (20 mM Tris/HCl, pH: 8) supplemented with cOmplete™ protease inhibitor (Roche) and 5 mM EDTA. Cell disruption was performed by mechanical impact force using accelerated glass beads (150-212 µm, Sigma) in a PreCellys homogenizer (Precellys Evolution, Bertin Technologies). Cell debris was pelleted via centrifugation. Remaining supernatant was further filtered using 0.45 and 0.2 µm membrane filters (Sarstedt). Filtered lysates with strep-tagged proteins were applied onto StrepTrap HP 1 ml columns (GE Healthcare). Subsequent protein separation was conducted using the ÄKTA start chromatography system (GE) according to the respective manufacturer’s protocols with following modifications: To prevent protein precipitation, 5 mM DTT were added to both the washing buffer (100 mM Tris/HCl, 150 mM NaCl, pH 8.0) as well as the elution buffer (100 mM Tris/HCl, 150 mM NaCl, 2.5 mM desthiobiotin, pH 8.0) for ctClpC purification. For bsMcsA, bsMcsB and gsMcsB, 10 mM DTT were added to the washing and the elution buffers. Buffer exchange was performed via centrifugal filters (Amicon® Ultracel®-30K, Merck) using buffer A supplemented with 30% glycerol and 5 mM DTT. Concentration and purity of desired proteins were determined via SDS-PAGE analyses followed by Bradford assays using a BSA standard curve.

### Arginine phosphorylation of β-casein

To obtain the ClpCP protease substrate β-casein-pArg, enzymatic arginine-phosphorylation of β- casein was performed in buffer BS (50 mM Tris/HCl [pH 8], 25 mM MgCl_2_, and 100 mM KCl, 2 mM DTT) in a total volume of 500 µl that was supplemented with an ATP regeneration system (2 mM ATP, 5 mM creatine phosphate, 2 U creatine phosphokinase). 5 µM arginine kinase (bsMcsAB or gsMcsB) and 50 µM β-casein were added accordingly. The phosphorylation reaction was incubated for 5 hours at 30 °C. Reverse purification using strep-tactin resin was employed to bind and remove bsMcsAB or gsMcsB from the flow-through. Anti-pArg antibodies were used to verify arginine phosphorylation of β-casein via dot blotting (Figure S1).

### *In vitro* casein degradation assays

Degradation of the protease substrate β-casein-pArg was performed in buffer PZ (25 mM HEPES, 200 mM KCl, 5 mM MgCl2, 1 mM DTT, 10% (v/v) glycerol, pH 7.6). Each reaction sample contained 2 µM bsMcsA, 2µM bsMcsB and 5 µM β-casein-pArg. 6 μM of total ctClpP in an equimolar ratio (total amount ctClpP1 and/or ctClpP2) and 2 µM ctClpC were added accordingly. Samples of 10 µl were taken from each reaction at specified time points during incubation at 32 °C. Substrate degradation was halted by adding sample loading buffer containing LDS (4x Bolt™ LDS Sample Buffer, Invitrogen) and heating at 80 °C. Degradation of the β-casein-pArg substrate was then analyzed via SDS-PAGE and subsequent Coomassie staining.

### *In vitro* ClpC basal ATPase activity assay

Quantitative determination of the basal ATPase activity of ClpC was performed using the ADP-Glo^TM^ system (Promega) in 384-well plates. ATPase reactions were carried out with 50 nM ultrapure ATP plus varying concentrations of ctClpC in buffer PZ. ATPase activity was determined according to the kit manual. Of note, unused ATP is depleted before ADP is converted to ATP for luciferase mediated luminescence quantification.

### *In vitro* ClpC chaperone activity assay

Quantitative analysis of ctClpC-mediated protection of luciferase aggregation during heat shock was adapted from Andersson et al. [78]. Luciferase from Photinus pyralis (0.2 µM) was heat-inactivated at 43 °C for 15 min in presence of ctClpC (0.4 µM) in buffer PZ or of bsClpC (0.4 µM) in buffer BS, both supplemented with 2 mM ATP. Adapter proteins (bsMcsA, bsMcsB and bsMecA) were added in an equimolar manner as indicated. β-casein (0.4 µM) as a substitute for ClpC served as negative control. After heat inactivation, all samples were immediately moved onto ice. 1 µl from each reaction sample was transferred into 199 µl buffer PZ (for ClpC) or buffer BS (for bsClpC) containing 400 µM D-luciferin and 20 mM ATP. Absolute luminescence was measured at the t = 5 s peak intensity.

### Determining protein-protein interactions with the BACTH system

To test protein-protein interaction between wild-type ClpC and mutants, we utilized the bacterial adenylate cyclase-based two-hybrid (BACTH) assay [50]. Genes to be examined are fused to one of either catalytic subunit, denoted as T18 and T25, of the *B. pertussis* adenylate cyclase. When the catalytic subunits are in close proximity, they can then reconstitute adenylate cyclase activity and allow growth of Δcya DHT1 *E. coli* on minimal medium with maltose. Both wild-type and mutant *clpC* genes were cloned into pKT25 or pUT18C vectors for testing of both homotypic and heterotypic interactions (see plasmid construction above). Each of the pKT25 and pUT18C plasmids were cotransformed into chemically competent DHT1 *E. coli* cells, and the cells were plated on a double antibiotic minimal M63 medium selection plate, which was supplemented with 0.5 mM IPTG for induction of the proteins, 40 µg/mL 5-bromo-4-chloro-3-indolyl-β-D-galactopyranoside (X-Gal), 0.04% casein hydrolysate, and 0.2% maltose. For a positive control, leucine zipper motifs were used in both pKT25 and pUT18C backgrounds with the appropriate antibiotic selection plates as they have previously been shown to interact [79]. Blue colonies indicating positive interactions were screened using the β-galactosidase assay. Positive colonies were chosen randomly and grown in M63 minimal medium with the appropriate antibiotics: 0.1% SDS and chloroform were then used to permeabilize the bacteria prior to addition of 0.1% o-nitrophenol-β-galactoside (ONPG), and 1 M NaHCO_3_ was used to halt the reaction after precisely 20 minutes of incubation time at room temperature. To quantify bacterial growth, an absorbance at 405 nm wavelength was recorded and normalized to bacterial growth (optical density of 600 nm [OD_600_]), the dilution factor, and the time (in minutes) of incubation prior to halting the reaction. The totals were reported in relative units (RU) of β-galactosidase activity.

### Purification of recombinant ClpC for antibody generation

The *clpC* gene from *C. trachomatis* L2 was cloned into pLATE31 (Invitrogen) to generate a C-terminal 6xHis-tagged construct. The vector was sequence verified and transformed into *E. coli* BL21 (DE3) *dAPX-1* for protein purification. Purification was performed using 2 L cultures based on the protocol described in reference [41]. Samples were induced with 100 mM isopropyl-β-D-thiogalactopyranoside (IPTG) and incubated with shaking for 20 h at 18 °C. Cultures were pelleted and frozen at −80 °C prior to purification. Samples were suspended in buffer A (25 mM Tris base [pH 7.5], 300 mM NaCl, and 20 mM imidazole), sonicated, ran through a 0.45 µm filter, and finally ran on an ÄKTA start chromatography system (GE). Sample was applied to a HisTrap™ FF 1 mL column. Proteins were eluted from the resin using buffer B (25 mM Tris base [pH 7.5], 300 mM NaCl, and 300 mM imidazole). Protein was concentrated and smaller proteins were removed using a Millipore Amicon Ultra 15 filtration unit (30 kDa cutoff). Protein was analyzed by SDS-PAGE to assess quality, then was purified further using a size exclusion chromatography (SEC) column on the ÄKTA. Purified protein from SEC was separated on SDS-PAGE to isolate appropriate fractions and was once again concentrated using a Millipore Amicon Ultra 15 filtration unit (30 kDa cutoff). Samples were sent to Thermo Fisher for polyclonal antibody generation in rabbits. Antibody specificity was determined by IFA and western blot comparing specific, and non-specific SEC fractions, and uninfected or infected McCoy Cells (see Fig. S2).

### Chlamydial transformation

The following protocol was a modification to the method developed by Mueller et al. [80]. For transformation, 10^6^ *C. trachomatis* serovar L2 EBs (25667R), naturally lacking the endogenous L2 plasmid, were incubated with 2 µg of plasmid in a volume of 50 µL CaCl_2_ at room temperature for 30 minutes. Each reaction was sufficient for a confluent monolayer of HeLa cells for a single well of a six-well plate, plated one day prior at 1 x 10^6^ cells. The 50 µL of transformant was added to a 1 mL overlay of room temperature Hanks’ balanced salt solution (HBSS) per well, followed by addition of 1 mL of HBSS per well. The six-well plate was centrifuged at 400 x *g* for 15 minutes at room temperature, and the beginning of this step was marked as the time of infection (0 hpi). Following centrifugation, the plate was incubated for 15 minutes at 37°C. At the end of the incubation period, the inoculum was aspirated and replaced with antibiotic-free DMEM containing 10 µg/mL gentamicin and 10% FBS. At 8 hpi, the medium was removed and replaced with DMEM containing 1 µg/mL cycloheximide, 10 µg/mL gentamicin, 1 U/mL penicillin, and 10% FBS. Cells infected with transformants were passaged every 48 hours by scraping cells from the plate and collecting the contents in a 2 mL microcentrifuge (mcf) tube. The mcf was then centrifuged at 17,000 x *g*, and the supernatant was aspirated, and the pellet resuspended in 1 mL HBSS. The resuspended pellet was then centrifuged again at 400 x *g* to remove host cell debris, and the inoculum was added dropwise to 1mL of HBSS on a new monolayer of HeLa cells. If a population of penicillin-resistant bacteria was established, then EBs were harvested by centrifuging scraped cells at 17,000 x *g*, aspirating supernatant, and resuspending in sucrose-phosphate (2SP) [66] and spun down at 400 x *g*. The EBs in the supernatant were then collected, and frozen at −80°C prior to stock titration. Plasmids were isolated from the strains using the DNeasy kit (Qiagen), re-transformed into *E. coli*, and re-isolated to verify insert by digest and sequencing.

### Western blot detection of endogenous HctB and ClpC or overexpressed ClpC_6xH

Crosslinking was performed for infections with an MOI = 1 with either wild-type Ctr L2, pBOMBL-ClpC_6xHis::L2, or pBOMBL-ClpC_Dbl_6xHis::L2 in a 6-well culturing plate. The pBOMBL-ClpC_6xHis::L2 and pBOMBL-ClpC_Dbl_6xHis::L2 transformants were induced at 16 hpi to reduce the drastic effects that were seen on inclusion morphology and bacteria number when induced at earlier time points. At time of sample collection, wells were washed three times with HBSS, and a separate well for IFA was fixed with MeOH as a control to monitor infection. Medium was aspirated and replaced with 1 mL HBSS pH 7.6 to better facilitate crosslinking. Disuccinimidyl suberate (DSS) in DMSO (100 mM) was added to crosslinked samples for a final concentration of 0.25 mM DSS. DMSO was added at the same volume to serve as vehicle control for non-crosslinked samples. 6-well plates were placed at 4°C on ice for at least 1 hour, aspirated, and 0.5 M Tris pH 7.6 was added to a final concentration of 20 mM per well for 10 minutes to quench the reaction. Samples were harvested in denaturing cell lysis buffer, quantified for protein using an EZQ Protein Quantification kit (Invitrogen), and separated on an 8% SDS-PAGE gel. Protein was transferred to PVDF membrane for western blot and stained with rabbit anti-ClpC antibody (for wild-type Ctr L2), rabbit anti-6xHis (for pBOMBL-ClpC_6xHis::L2 and pBOMBL-ClpC_Dbl_6xHis::L2 transformants; Abcam), or rabbit anti-HctB (for wild-type Ctr_L2; kind gift of Dr. T. Hackstadt) and goat anti-MOMP of *Chlamydia* (Meridian Bioscience). Donkey secondary antibodies conjugated to fluorophores were used to detect rabbit (680 nm) and goat (800 nm) primary antibodies (Licor). Labeling was observed on an Azure Biosystems c600 instrument using the NIR autoexposure setting.

### Determining the effect of overexpression of wild-type and mutant ClpC proteins or knockdown of *clpC* via immunofluorescence and inclusion-forming unit analysis

*C. trachomatis* transformed with either the pBOMBL-ClpC_6xHis::L2 isoforms, the dual expression plasmid pBOMBL-ClpC_6xHis-ClpX_FLAG, or pBOMBL12CRia(*clpC*) under the control of an aTc-inducible promoter were used to infect a monolayer of HeLa cells on coverslips utilizing penicillin as a selection agent. Samples were induced with increasing concentrations of aTc at either 4 or 10 hpi and were fixed with methanol after a 20 h or 14 h pulse (24 hpi). Fixed cells were incubated with goat anti-MOMP primary antibody (all IFA samples), mouse anti-6xHis (overexpression plasmids), rabbit anti-FLAG (pBOMBL-ClpC_6xHis-ClpX_FLAG only; Sigma-Millipore), or mouse anti-Cpf1 (pBOMBL12CRia(*clpC*); Sigma-Millipore). Donkey anti-goat Alexa Fluor 594-conjugated secondary antibody were used for visualization of the organisms for experiments. A donkey anti-mouse Alexa Fluor 488-conjugated secondary antibody was used for visualization of ClpC and isoforms as well as dCas12, respectively, for expression and localization. To visualize ClpX_FLAG, a donkey anti-rabbit Alexa Fluor 647-conjugated secondary antibody was used to visualize expression and localization. Lastly, samples were stained with 4’,6-diamidino-2-phenylindole (DAPI) to visualize the host and bacterial cell DNA. Representative images were taken on a Zeiss LSM 800 confocal laser scanning microscope with a 100x objective, then 5.5x digitally zoomed and adjusted for color and brightness with Zen software (blue edition, V3.3).

To assess the effects of wild-type and ClpC mutant overexpression, the inclusion forming unit (IFU) assay was used. HeLa cells were infected in triplicate as described above for IFA samples. IFUs were harvested by scraping sample wells in 2SP, pooling triplicate sample wells together, and freezing at least overnight at −80°C. Samples were then removed, vortexed, and serially diluted in HBSS and titrated in duplicate directly onto a new monolayer of HeLa cells. Following 24-40 hours of incubation, the samples were fixed and stained with goat anti-MOMP primary antibody, and a donkey anti-goat Alexa Fluor 594-conjugated secondary antibody for visualization of the bacteria within inclusions. An aldehyde fix was used if plasmid retention was to be observed, to retain GFP fluorescence, followed by MeOH permeabilization. Ten fields of view were counted for each duplicate well at 20x magnification, giving a total of 20 fields of view per experiment. Three independent experiments were performed, and the totals for each experiment were averaged. Displayed values were expressed as a percentage of the uninduced sample to provide an internal control. For statistics, a parametric, unpaired Student’s two-tailed *t* test was used to compare the induced samples to the uninduced control and was performed using the averages of each biological replicate.

### Knockdown of endogenous *C. trachomatis* ClpC

Two wells of a six well plate per condition were infected using *C. trachomatis* transformed with pBOMBL12CRia(*clpC*)::L2 at an MOI of 1.0. At 10 hpi samples were induced or not with 2 nM aTc. For each designated time point, RNA was collected using TRIzol reagent (Invitrogen) and extracted using chloroform with the aqueous phase precipitated with an equal part of isopropanol, according to the manufacturer’s instructions (Thermo Fisher). DNA was removed from RNA samples through rigorous DNA-free treatment (Thermo Fisher) before 1 µg was reverse transcribed with Superscript III reverse transcriptase (RT; Thermo Fisher). Equal volumes of cDNA were used for qPCR. For genomic DNA, cells were trypsinized and collected by centrifugation for 5 minutes at 400 x g and resuspended in phosphate-buffered saline (PBS) then stored at −20°C until further processing. Frozen samples were thawed and subjected to a DNeasy Blood and Tissue kit (Qiagen) to isolate *C. trachomatis* DNA according to the manufacturer’s instructions. 5 ng/µL was used for qPCR. Transcripts and genomic DNA were quantified by qPCR in 25 µL reaction mixtures using 2x SYBR green master mix in an ABI700 thermal cycler in comparison to a standard curve generated from purified *C. trachomatis* L2 genomic DNA. Transcripts were then normalized to genomic DNA for results and the induced samples normalized to the uninduced samples for the same time points.

## Acknowledgements

We would like to thank Drs. H. Caldwell (NIH/NIAID) for eukaryotic cell lines, T. Hackstadt (RML/NIAID) for the pBOMB4-Tet::L2 plasmid and the rabbit anti-HctB antibody, I. Clarke (University of Southampton) for the plasmidless strain of *C. trachomatis* serovar L2, and Mr. Lucas Struble in Dr. G. Borgstahl’s lab (UNMC) for his help purifying recombinant ClpC protein for antibody production. We further thank Drs. Paul R. Thompson (UMass Chan Medical School) for providing anti-P-Arg antibodies and Tim Clausen (IMP Vienna) for sharing the plasmid encoding *G. stearothermophilus* McsB. This project was supported by an NIAID/National Institutes of Health awards (1R56AI146062 to S.P.O. and 1R01AI170688 to S.P.O and D.J.F) and university funds to D.J.F. We further appreciate funding by the Deutsche Forschungsgemeinschaft (DFG, German Research Foundation): TRR 261, Project-ID 398967434 to P.S., H.B.O., and B.H., as well as support by infrastructural funding from Cluster of Excellence EXC 2124 Controlling Microbes to Fight Infections, Project-ID 390838134.

---

**Supplemental Figure 1:**
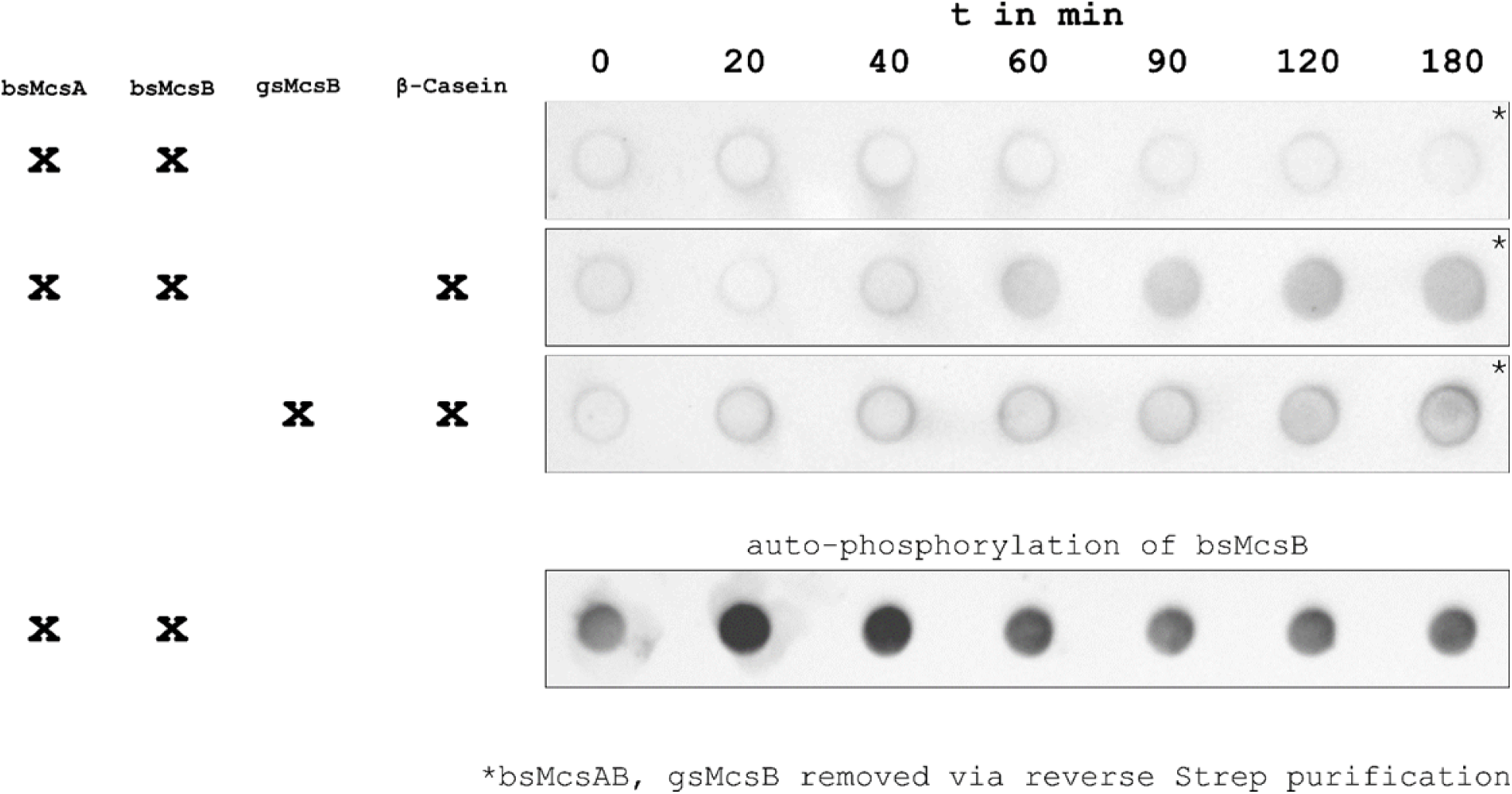
McsB mediated arginine phosphorylation of β-casein. Dot-blot analysis using anti-pArg-antibodies. Strep-tagged McsAB of *B. subtilis* (bsMcsAB) or McsB of *G. stearothermophilus* (gsMcsB) were removed prior via reverse strep purification as indicated by *. β-casein is gradually arginine-phosphorylated by bsMcsAB or gsMcsB. In the reverse purified bsMcsAB fraction, auto-phosphorylation of presumably bsMcsB occurs rapidly.

**Supplemental Figure 2:**
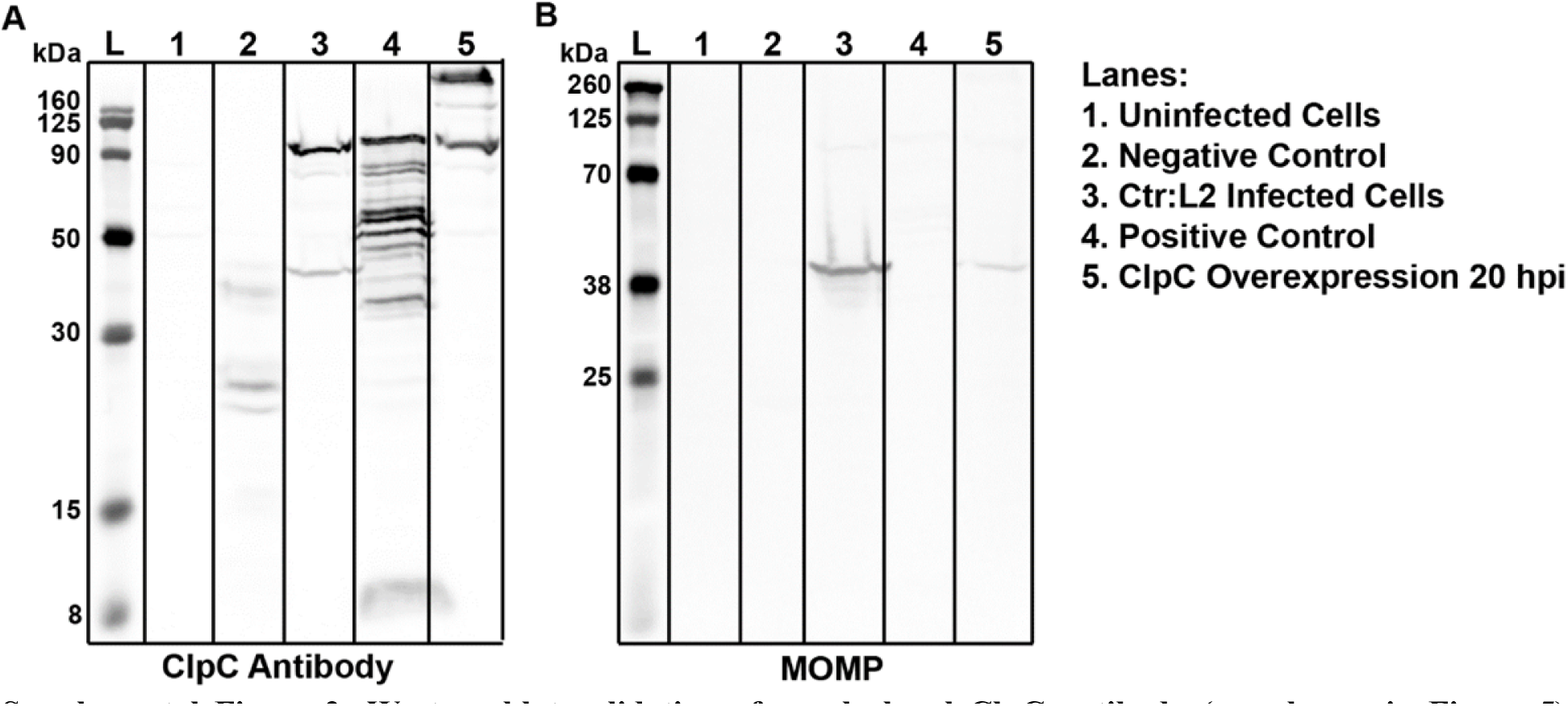
Western blot validation of a polyclonal ClpC antibody (see also main Figure 5). Western blot stained with primary antibodies of either (A) a ClpC antibody from rabbit serum, or (B) MOMP for confirmation of Ctr infection. ClpC = 95.2 kDa, MOMP = 42 kDa. Lanes 1 and 3 are whole cell lysates collected from McCoy cells either uninfected or infected with a wild-type *C. trachomatis* serovar L2 Bu/434. The negative and positive controls are size exclusion chromatography fractions for nonspecific or ClpC specific fractions, respectively. Multiple bands in lane 4 are likely a result of ClpC degradation over time. Lane 5 was collected from HeLa cells infected with Ctr:pBOMBL(ClpC), crosslinked at the time indicated, and performed as described in material and methods. Antibody was used 1:2000 in 1% PBST as described in the materials and methods.

**Supplemental Figure 3:**
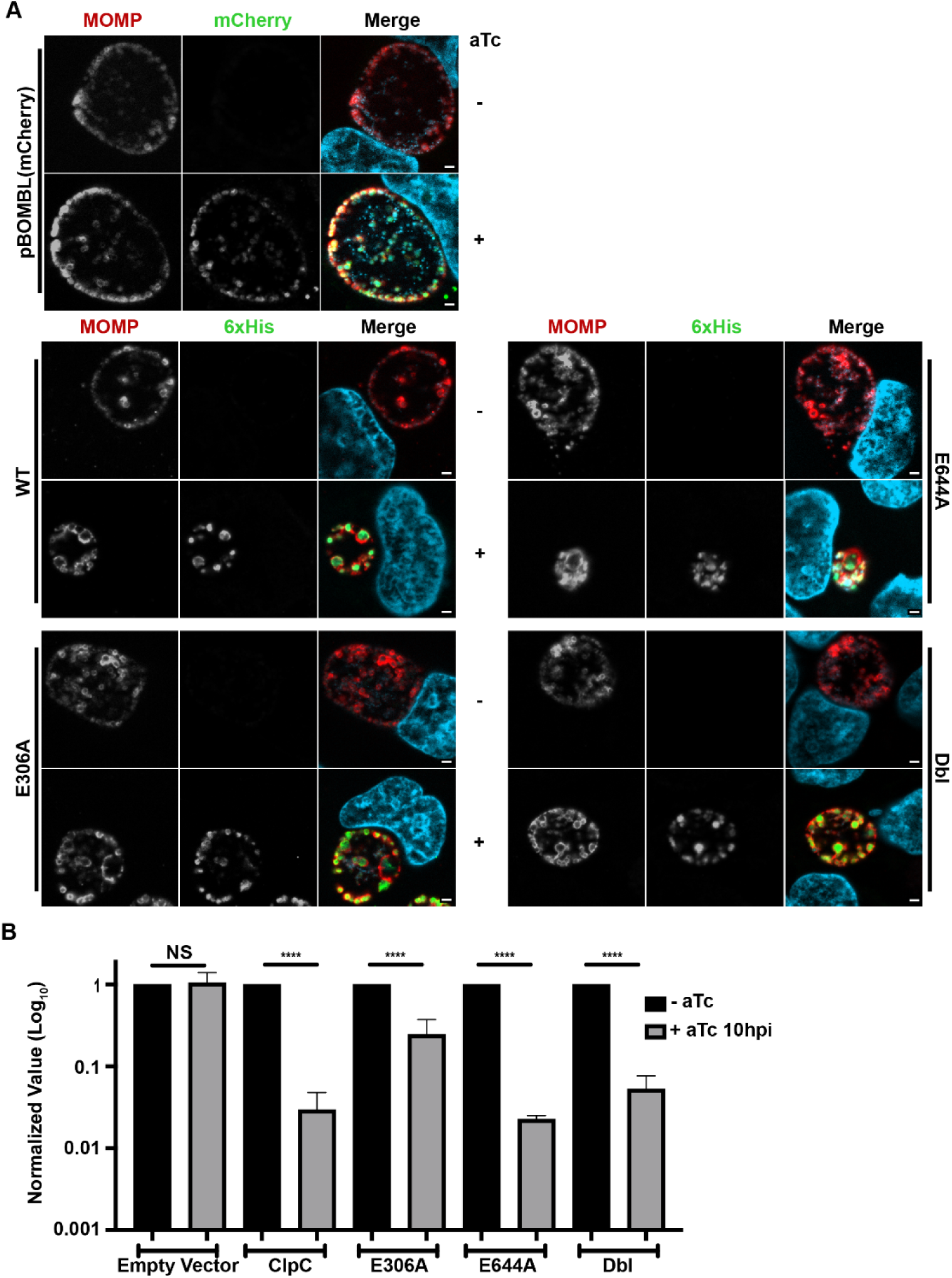
Overexpression of ClpC isoforms shows NBD1 is critical for function even when expressed at 10 h post-infection (see also main Figure 6). (A) Immunofluorescence analysis of the control plasmid expressing mCherry, or exogenously overexpressed ClpC isoforms as indicated with C-terminal 6xHis tags induced at 10 hpi with 20 nM aTc. At 24 hpi the control plasmid was fixed with 3.25% formaldehyde and 0.125% glutaraldehyde followed by MeOH to retain mCherry fluorescence, while the ClpC isoforms were fixed with MeOH. Samples were stained for MOMP (green/red) or 6xHis (green) tagged protein from plasmid induction, and DNA (light blue). Representative images were taken in triplicate on a Zeiss Apotome at ×100 magnification with ×5.5 digital zoom. Scale bar = 2 µm. (B) Recovered IFUs of cells infected with each transformant from (A). Samples were collected in triplicate and combined at 24 hpi, frozen at −80 °C, and titrated onto a fresh monolayer of cells to quantify inclusions in the secondary infection. Inclusions were identified by staining with MOMP and a secondary antibody conjugated to a fluorophore. Data are the average of at least three independent experiments. The statistical significance analysis was performed via a parametric, unpaired Students T-test (NS = not significant; ****p<0.0001).

**Supplementary Figure 4:**
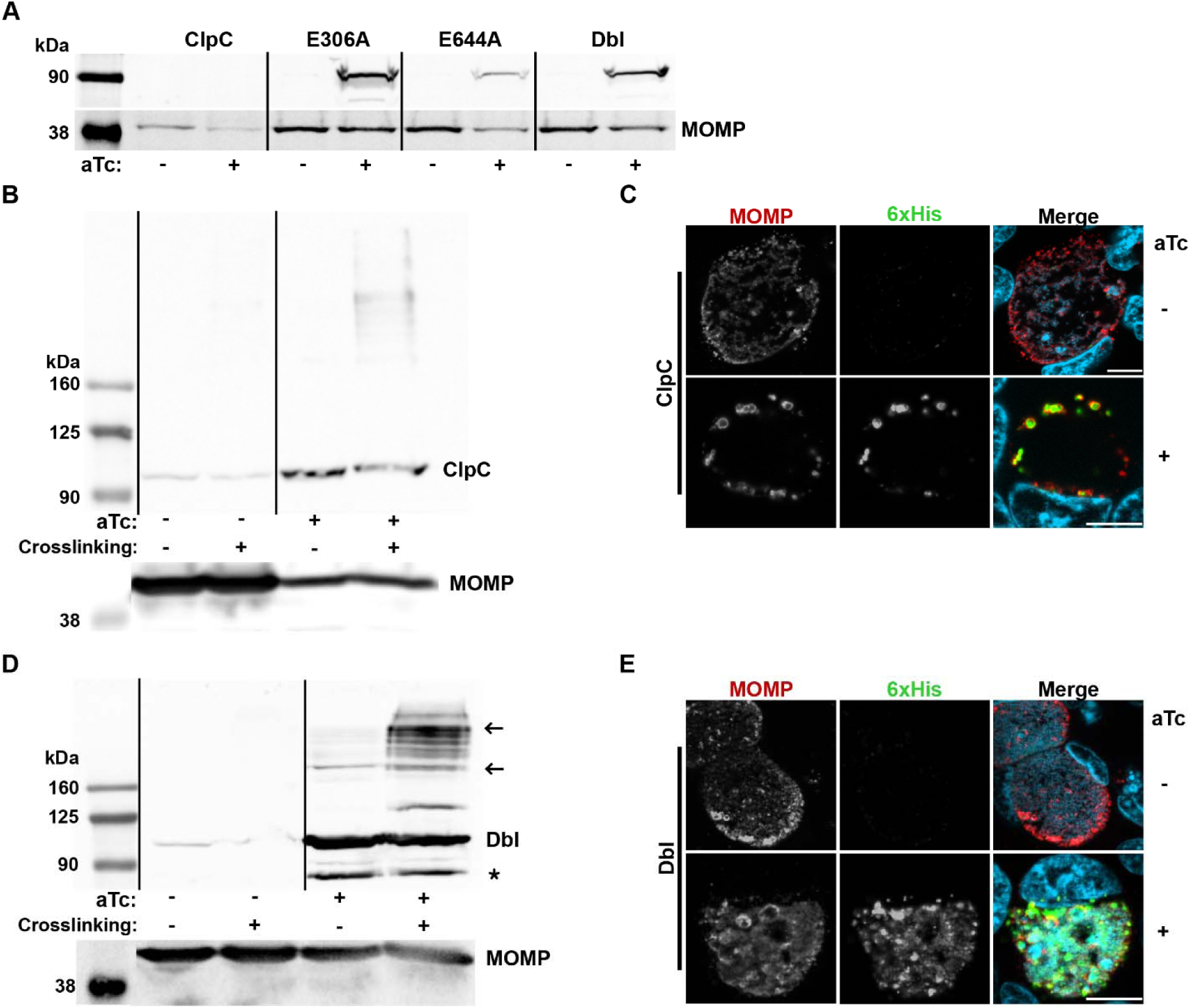
Expression levels and oligomerization states of ClpC_6xHis isoforms *in vivo* under different conditions. (A) Detection of ClpC_6xHis isoforms in infected cells. Whole cell lysates were collected at 24 hpi from cells infected with the indicated ClpC_6xHis isoform after inducing expression or not with 20 nM aTc at 4 hpi. Samples were processed for western blotting using 45-50 µg of protein per lane, and the ClpC isoform was detected using an anti-6xHis antibody. See also main Figure 6. (B-E) Exogenously expressed ClpC_6xHis isoforms can oligomerize *in vivo*. Western blots of whole cell lysates for the (B) wild-type ClpC_6xHis and (D) Dbl mutant ClpC_6xHis isoform and (C,E) IFA controls of overexpressed (C) ClpC_6xHis or (E) Dbl mutant ClpC_6xHis. Whole cell lysates for (B,D) were collected after inducing expression or not with 20 nM aTc at 16 hpi and collected at 48 hpi. Some samples were crosslinked as indicated prior to harvesting lysates. 25-50 µg of protein from samples were loaded per lane. Arrows indicate predicted dimer or hexamer band, with the asterisk indicating a commonly seen band when staining for ClpC. See also main Figure 5. ClpC = 95.2 kDa, MOMP = 42 kDa. C and E, Scale bar = 10 µm.

**Supplementary Figure 5:**
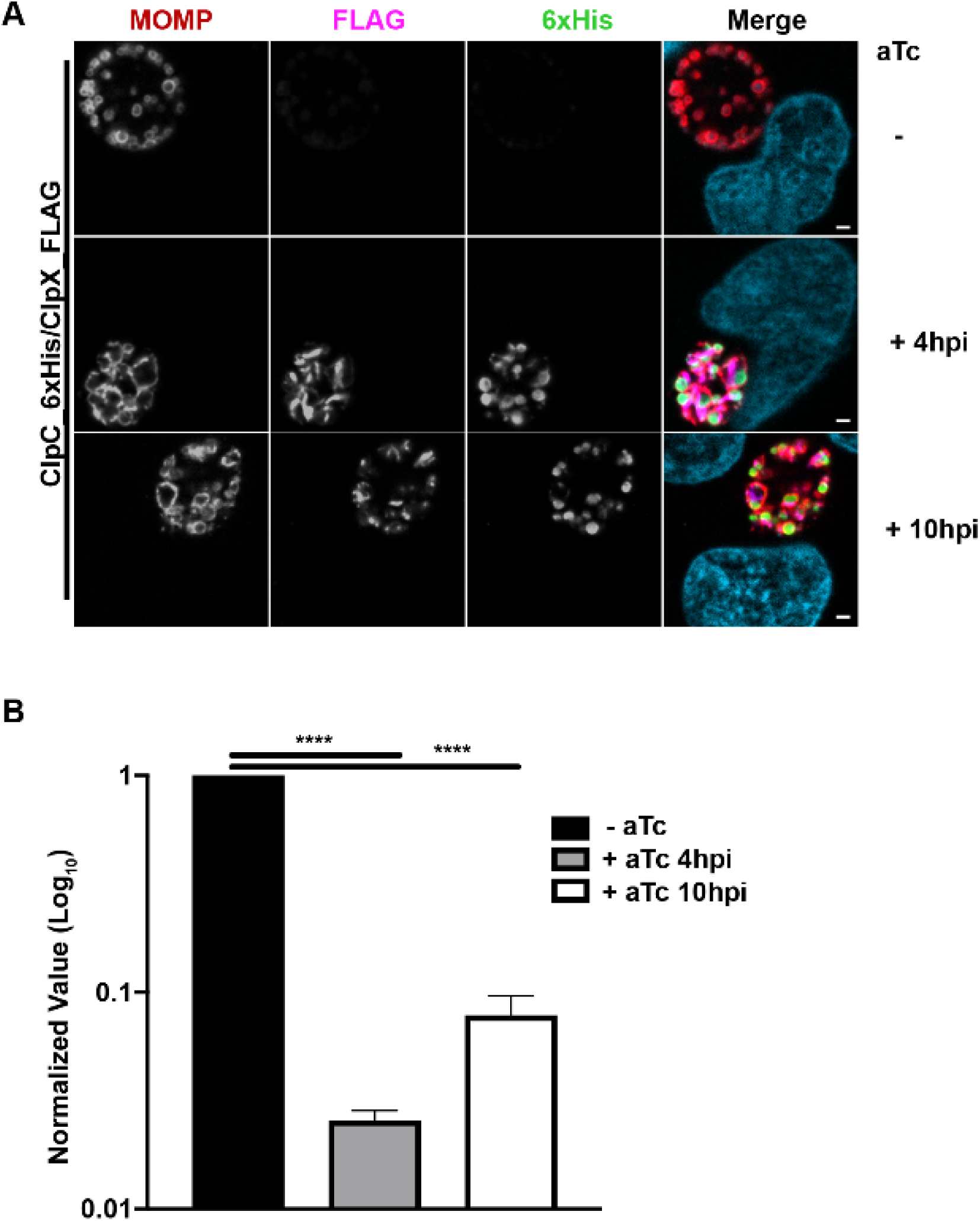
Overexpression of ClpC is not a result of titrating ClpP away from ClpX. (A) Immunofluorescence analysis of exogenous dual expression of ClpC_6xHis and ClpX_FLAG induced at 4 or 10 hpi with 20 nM aTc and fixed at 24 hpi with MeOH. Samples were stained for MOMP (red), 6xHis (green - ClpC) or FLAG (pink - ClpX) tagged protein from plasmid induction, and DNA (light blue). Representative images were taken in triplicate on a Zeiss Apotome at ×100 magnification with ×5.5 digital zoom. Scale bar = 2 µm. (B) Recovered IFUs of cells infected with each transformant from (A). Samples were collected in triplicate and combined at 24 hpi, frozen at −80 °C, and titrated onto a fresh monolayer of cells to quantify inclusions in the secondary infection. Inclusions were identified by staining with MOMP and a secondary antibody conjugated to a fluorophore. Data are the average of three independent experiments. The statistical significance analysis was performed via a parametric, unpaired Students T-test (****p<0.0001). See also main Figure 6 for comparison.

**Supplementary table 1:**
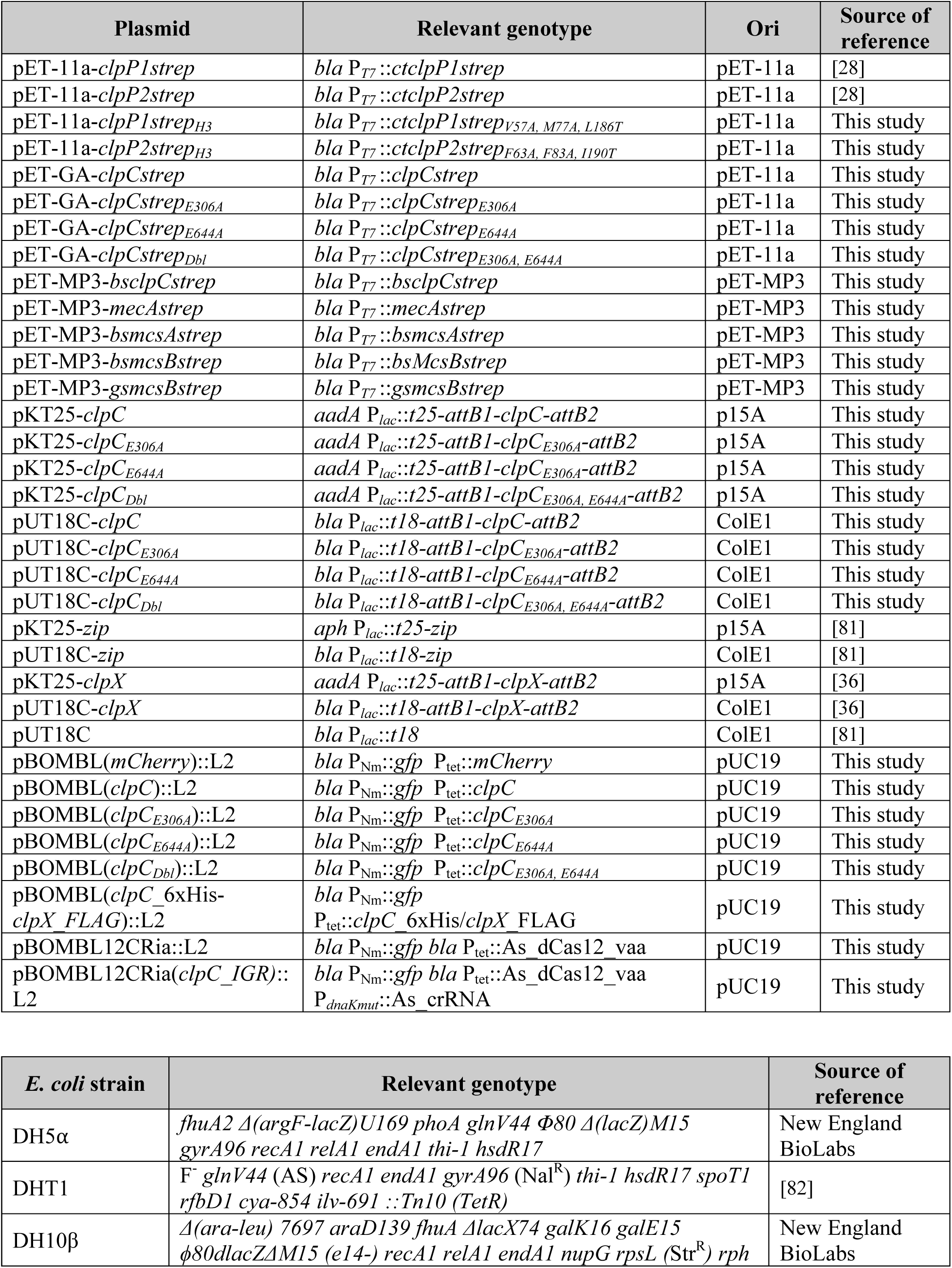
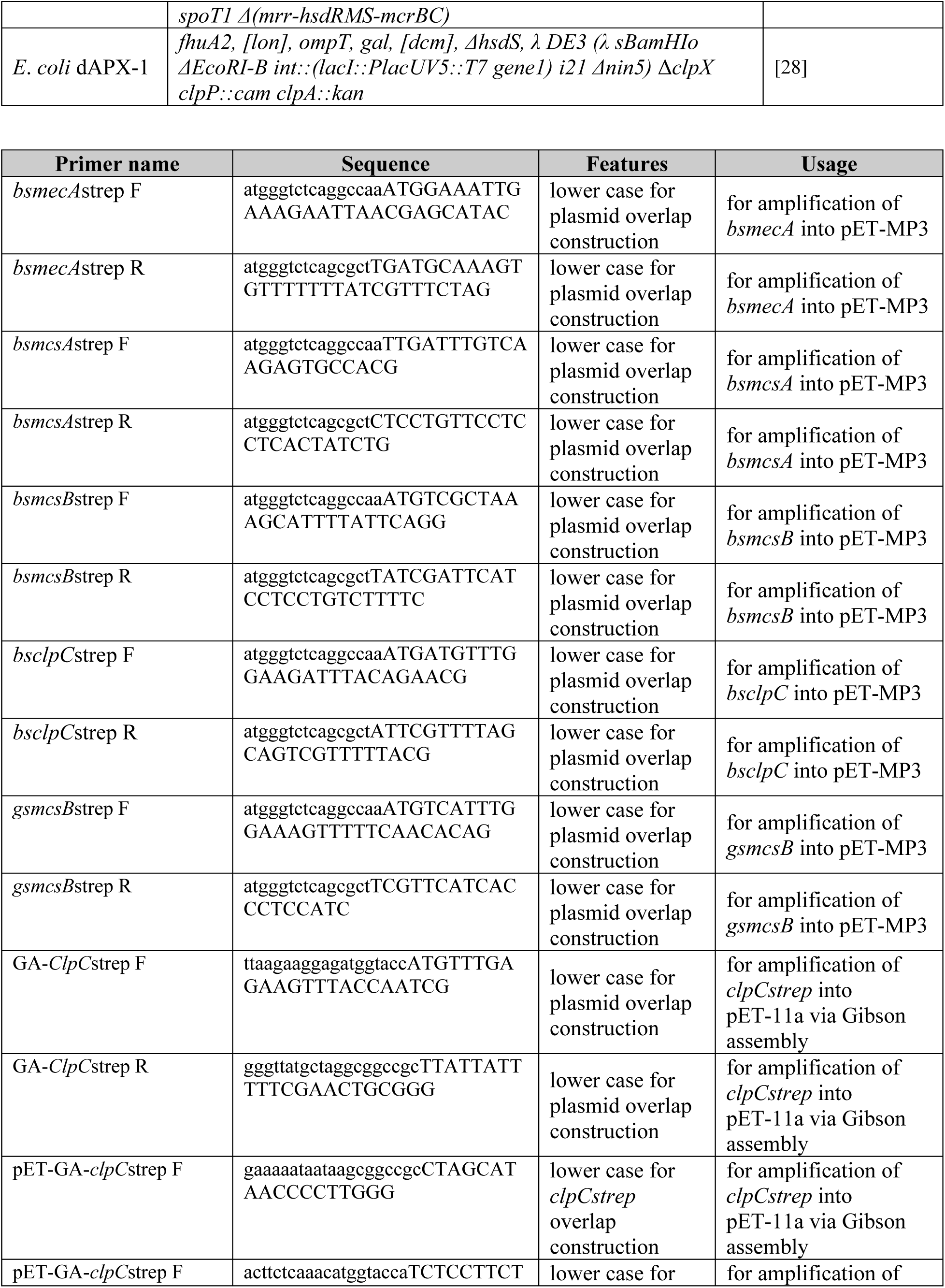
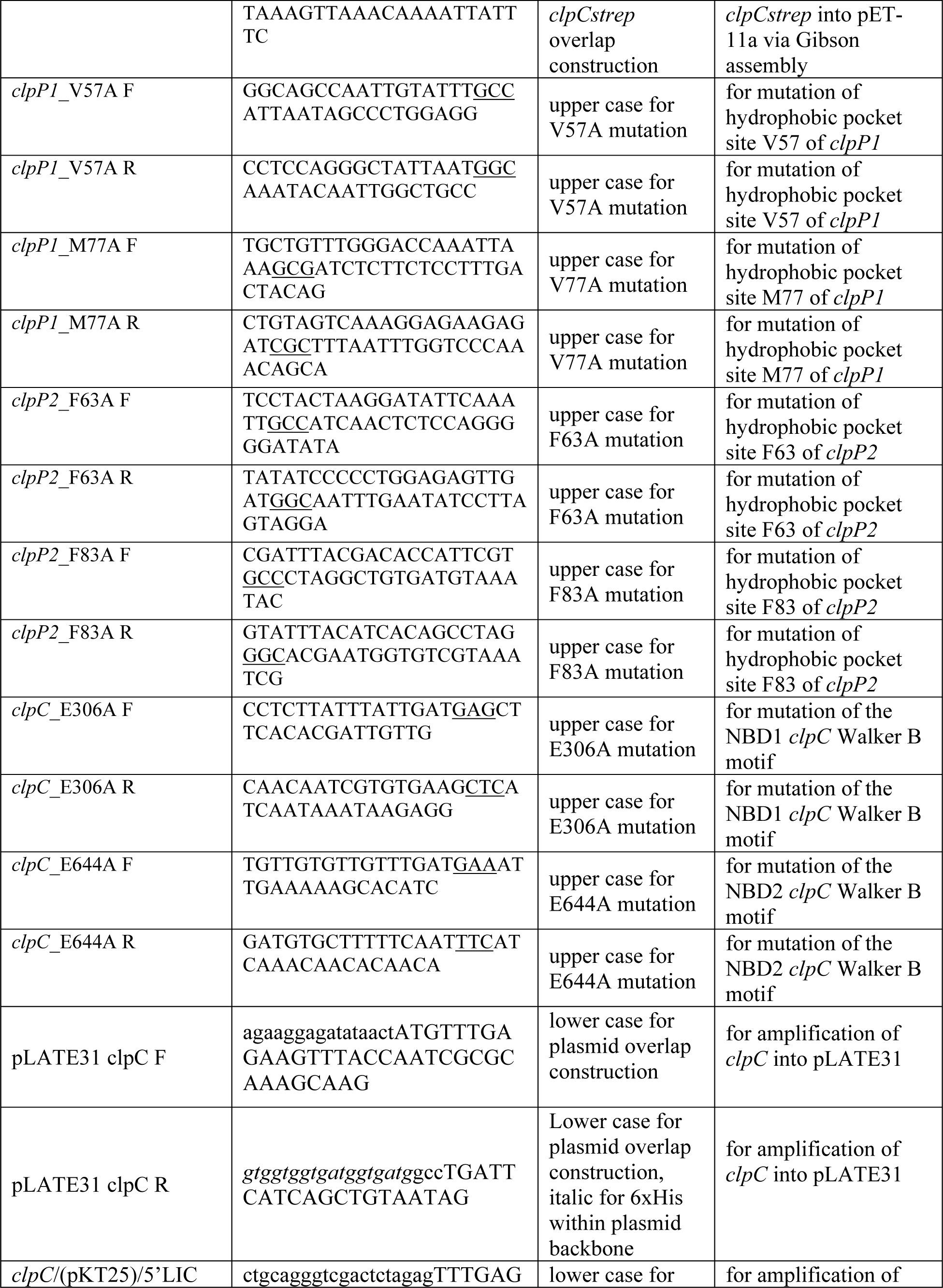
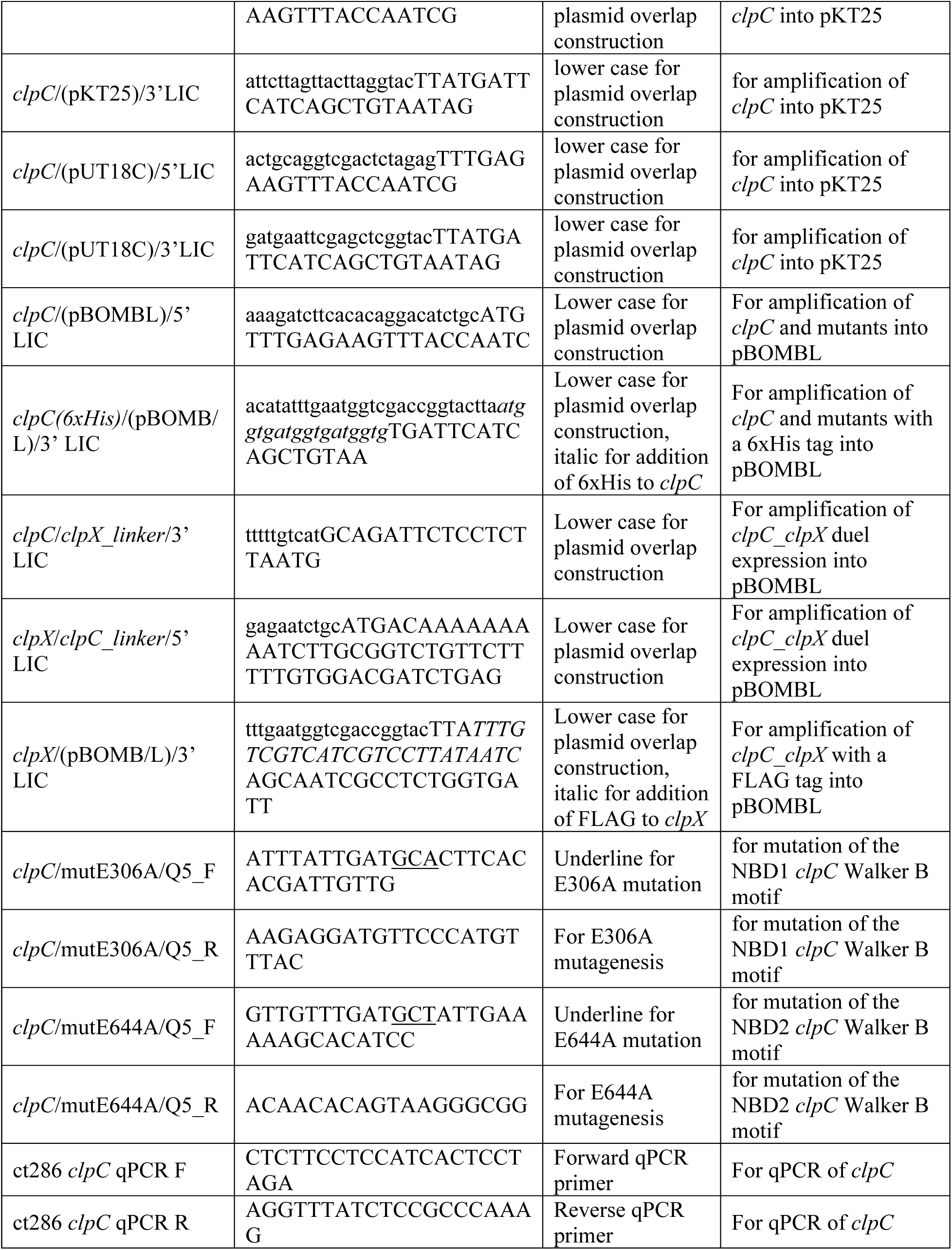
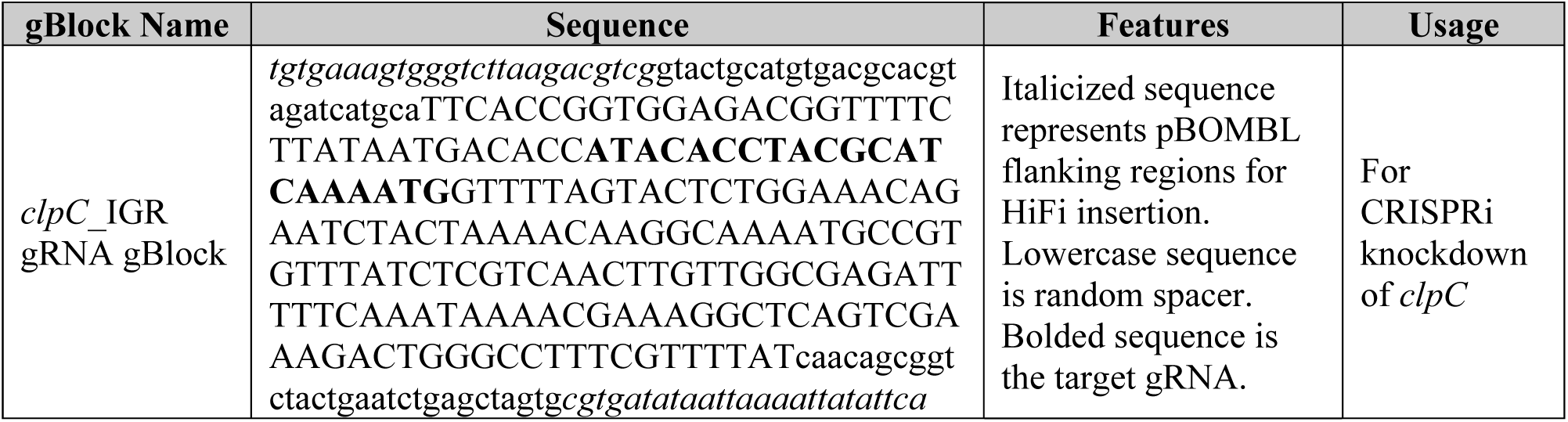
List of Plasmids, Strains, and Primers

